# NR4A1 limits CD8^+^ T Cell effector responses and protection in tuberculosis

**DOI:** 10.64898/2026.01.12.699051

**Authors:** Samreen Fatima, Yao Chen, Lorissa Smulan, Basanthi Satish, Mike Jameson, Sheetal Saini, Calvin Johnson, Alicia Tay, Shihui Foo, Hedayathulla Rahmathulla, Akhila Balachander, Shanshan W. Howland, Hardy Kornfeld, Amit Singhal

## Abstract

During *Mycobacterium tuberculosis* (*Mtb*) infection, CD8^+^ T cells exhibit dysfunction with impaired cytotoxicity and limited localization to granuloma cores. Using knockout mice, adoptive-transfer models and validation in macaque and human datasets, we identified the nuclear receptor NR4A1 as a key restrainer of CD8^+^ T cell immunity in tuberculosis (TB). *Mtb*-infected *Nr4a1*^-/-^ mice displayed reduced bacterial burden, attenuated pathology, higher lung CD8^+^/CD4^+^ T cell ratios, and enhanced CD8^+^ T cell effector functions. Bulk and single-cell RNA sequencing revealed suppression of gene expression program linked with exhaustion, and expansion of *Nkg7*^+^ and *Granzyme*^+^ cytotoxic CD8^+^ T cell subsets in *Nr4a1*^-/-^ mice. Spatial analyses demonstrated increased infiltration of *Nkg7*^+^ activated CD8^+^ T cells in *Nr4a1*^-/-^ lesions. ChIP-qPCR showed NR4A1 binding to *Nkg7* promoter, and *Nkg7* knockdown abrogated the enhanced cytotoxicity of *Nr4a1*^-/-^ CD8^+^ T cells. Pharmacologic inhibition of NR4A1 reduced *Mtb* burden and pathology, and restored *Nkg7* expression and CD8^+^ T cell infiltration in the lung. Together, these findings identify NR4A1 as a negative regulator of CD8^+^ T cell–mediated immunity in TB and suggest the NR4A1-NKG7 axis as a novel host-directed therapeutic target.

**A one-sentence summary of your paper:** NR4A1 suppresses CD8^+^ T cell infiltration and cytotoxicity in TB lesions, and its inhibition enhances host resistance to *Mtb* infection.

## Introduction

Tuberculosis (TB), caused by *Mycobacterium tuberculosis* (*Mtb*), remains the world’s deadliest infectious disease, with an estimated 10.8 million new cases and 1.25 million deaths in 2023^1^. Despite available antibiotics, TB continues to drive global mortality, particularly among individuals living with HIV and those infected with drug-resistant *Mtb* strains. Control of *Mtb* infection depends upon the coordinated expression of adaptive immunity. CD4^+^ T cells play the major role in TB immunity by producing cytokines like IFN-γ and TNF-α that help infected macrophages and restrict bacterial growth^2^. While CD8^+^ T cells can contribute to control of TB through cytotoxic and cytokine functions, their relative importance in vivo remains debated^3, 4, 5^. Importantly, CD8^+^ T cells are spatially excluded from granuloma cores-primary sites of active bacterial replication,^7, 8^ limiting their access to infected macrophages^9^. During chronic *Mtb* infection they also exhibit poor cytotoxic differentiation and show signs of functional exhaustion^10, 11^.

The orphan nuclear receptor NR4A1 (also known as Nur77) is an immediate-early gene known to regulate gene expression important for T cell exhaustion, activation and tolerance^12^. In chronic infections and cancer, NR4A1 is strongly induced in antigen-stimulated T cells, where it acts as a transcriptional repressor of key effector programs. It competes with AP-1 at regulatory DNA sites, dampening expression of cytotoxic mediators like IFN-γ and granzymes while favoring the maintenance of progenitor-exhausted subsets^13^. Beyond exhaustion, NR4A1 also contributes to peripheral tolerance and anergy by regulating *Egr2*, *Ctla4*, and *Pdcd1*, acting as a key transcriptional switch that limits T cell–mediated inflammation^14, 15, 16^. Given the persistent antigen stimulation and functional exhaustion characteristic of TB, we hypothesized that NR4A1 acts as a molecular brake on CD8^+^ T cell-mediated protective immunity.

In this study, we investigated the regulatory role of NR4A1 in *Mtb* infection and disease progression using murine models, with validation in macaque and human datasets. Genetic deletion and pharmacological inhibition of NR4A1 significantly improved resistance to TB, primarily through heightened cytotoxic potential and reduced exhaustion of CD8^+^ T cells. We showed that NR4A1 binds to the *Nkg7* promoter, repressing transcription. NR4A1 deficiency derepresses NKG7 expression and facilitates infiltration of CD8^+^ T cells into TB lesions. These findings establish NR4A1 as a transcriptional checkpoint and a critical negative regulator of CD8^+^ T cell-mediated protective immunity, highlighting its potential as a target for host-directed therapy.

## Results

### *Nr4a1* deletion enhances resistance to *Mtb* in mice

Wild type (WT) and *Nr4a1*^-/-^ (KO) mice were infected with a low aerosol dose of *Mtb* Erdman (Fig. 1a). In *Mtb-*infected WT mice, *Nr4a1* mRNA levels in the lung increased by 2-fold at 8-weeks (wk) post-infection (Extended Data Fig. 1a). Remarkably, *Nr4a1*^-/-^ mice demonstrated a significant survival advantage compared to WT (Fig. 1b). Compared to WT mice, *Nr4a1*^⁻/⁻^ mice consistently exhibited significantly lower *Mtb* colony-forming units (CFU) in lung and spleen at 4, 8, and 16 wk (Fig. 1c–e). Analysis of H&E-stained lung tissue sections revealed striking differences between the genotypes. While WT mice developed large, well-structured lung lesions, lung sections from *Nr4a1*^⁻/⁻^ mice showed smaller, more diffuse infiltrates (Fig. 1f). Quantitative morphometry confirmed significantly reduced immune pathology in *Nr4a1*^⁻/⁻^ lungs at all time points (Fig. 1g). These data indicated that NR4A1 limits host control of *Mtb*, while its absence facilitates a more effective host response.

**Fig. 1.**
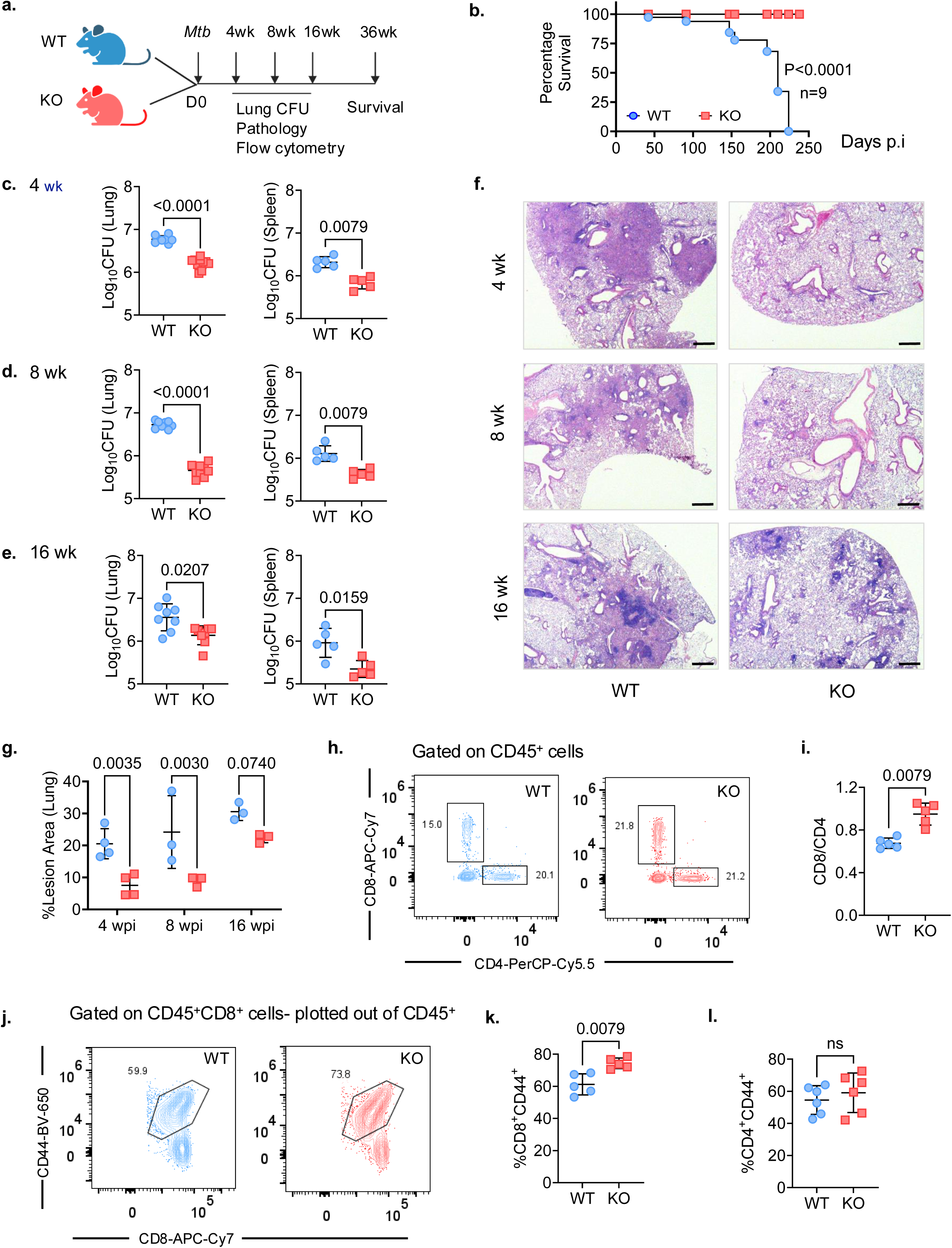
Deletion of NR4A1 results in enhanced host resistance to *Mtb* in murine model. **a**, Schematic model of the growth kinetics and survival experiment. **b**, Percent survival over time in days for wildtype (WT) and *Nr4a1*^-/-^ (KO) mice (*n* = 9 per group) (*P* < 0.0001, Mantel–Cox test). **c-e**, Bacterial load in lungs—left panel and spleen—right panel in WT versus *Nr4a1*^−/−^ mice infected with *Mtb* by aerosol and euthanized at 4, 8 and 16 wk. **f**, Representative images of H&E (purple) stained lung sections from *Nr4a1*^−/−^ (right) and WT mice (left) at the indicated time points after infection. Each image is representative of tissue sections from at least 3 individual mice from 2 experiments (scale bar=100 µm). **g**, TB lesion area expressed as percentage of total lung area in WT (Blue circles) and *Nr4a1^-^*^/-^ mice (red squares) at 4, 8 and 16 wk. *P* values were determined using two-tailed, Mann–Whitney test. **h**, Representative Flow-cytometric plots of CD8^+^ versus CD4^+^ T cells among CD45^+^ cells isolated from the lungs of WT (blue) and *Nr4a1*^−/−^ (red) at 8 wk *(n* = 5 mice per group). **i**, Ratio of CD8^+^ to CD4^+^ cells among CD45^+^ cells isolated from the lungs of WT (blue) and *Nr4a1*^-/-^ mice (red). **j,k**, Representative FACS plots (j) and quantification (k) of activated (CD44^+^) CD8^+^ cells among lung CD45^+^ cells in WT and *Nr4a1*^−/−^ mice at 8 wk. **l**, quantification of activated (CD44^+^) CD4^+^ cells among lung CD45^+^ cells in WT and *Nr4a1*^−/−^ mice at 8 wk. Data in c-e, i, k and l as mean ± s.d. representative of 2-3 independent experiments. *P values*, two-tailed Mann–Whitney test. ns, not significant.

Since NR4A1 is highly expressed in activated T cells and regulates T cell exhaustion and activation thresholds^16, 17, 18, 19, 20^, we evaluated the T cell compartment in infected lungs (Extended Data Fig.1b). Flow-cytometric analysis at 8 wk revealed a significantly increased ratio of CD8^+^ to CD4^+^ T cells in *Nr4a1*^⁻/⁻^ lungs compared to WT (Fig. 1h,i). Lung CD8^+^ T cells in *Nr4a1*^⁻/⁻^ mice also showed a higher frequency of CD44^+^ activated cells (Fig. 1j,k). No change in the activation status of the CD4^+^ T cells was observed (Fig. 1l). These results suggested that NR4A1 affects CD8^+^ T cell activation and expansion during *Mtb* infection. Noteworthy, the composition of lung myeloid cells among *Mtb*-infected WT and *Nr4a1*^⁻/⁻^ mice were similar, except for a modest reduction in neutrophil percentage in *Nr4a1*^⁻/⁻^ lungs (Extended Data Fig. 1c-g). Collectively, these findings demonstrated that genetic deletion of NR4A1 enhances immune control of *Mtb* infection, identifying NR4A1 as a negative regulator of protective immunity in TB.

### Distinct gene expression signatures among *Nr4a1*^⁻/⁻^ CD8^+^ T cells

NR4A1 can both activate and repress gene expression^13, 18, 21, 22^. To characterize how NR4A1 shapes T cell transcriptional programs during *Mtb* infection, we performed a time-course bulk RNA-seq of magnetically sorted splenic CD8^+^ T cells from *Mtb*-infected and uninfected (UI) WT and *Nr4a1*^⁻/⁻^ mice. Principal Component analysis (PCA) revealed distinct transcriptional profiles by infection status and genotype (Fig. 2a). Time course analysis identified 463 differentially expressed genes (DEGs), forming seven clusters with distinct temporal patterns (Fig. 2b,c, Supplementary Table 1). Clusters 5 and 6 were enriched in UI *Nr4a1*^⁻/⁻^ CD8^+^ T cells compared to UI WT CD8^+^ T cells (KO-UI vs WT-UI, Fig. 2b). Cluster 5 genes encoded proteins related to leukocyte migration (*Ccr1*, *Cxcr2*, *Trem1*, *Fcer1g*, *Ccl6*, *Ccl24*) and cytokine-mediated signaling (*Il36g*, *Il1b*, *Il1rn*, *Cd44*, *Tnfsf18*); cluster 6 genes encoded antibacterial defense (*Ltf*, *Ncf1*, *S100a8*, *Fosl2*) and leukocyte migration proteins (*Cd9*, *Cxcl2*, *Anxa1*). Most of these support CD8^+^ T cell activation, differentiation, and cytotoxicity. The temporal trajectories of clusters 1, 3, 4 and 7 were significantly different among WT and *Nr4a1*^⁻/⁻^ CD8^+^ T cells over infection (2 – 8 wk; Fig. 2c). Cluster 1 genes were consistently upregulated in *Nr4a1*^⁻/⁻^ CD8^+^ T cells and included *Pbx4* and *Slc1a5*, which correlates with the CD8^+^ T cell infiltration^23, 24^. Cluster 4 genes were downregulated and included *Ubd* and *Dhdh*, reported to impair T cell proliferation and function^25, 26^.

**Fig. 2.**
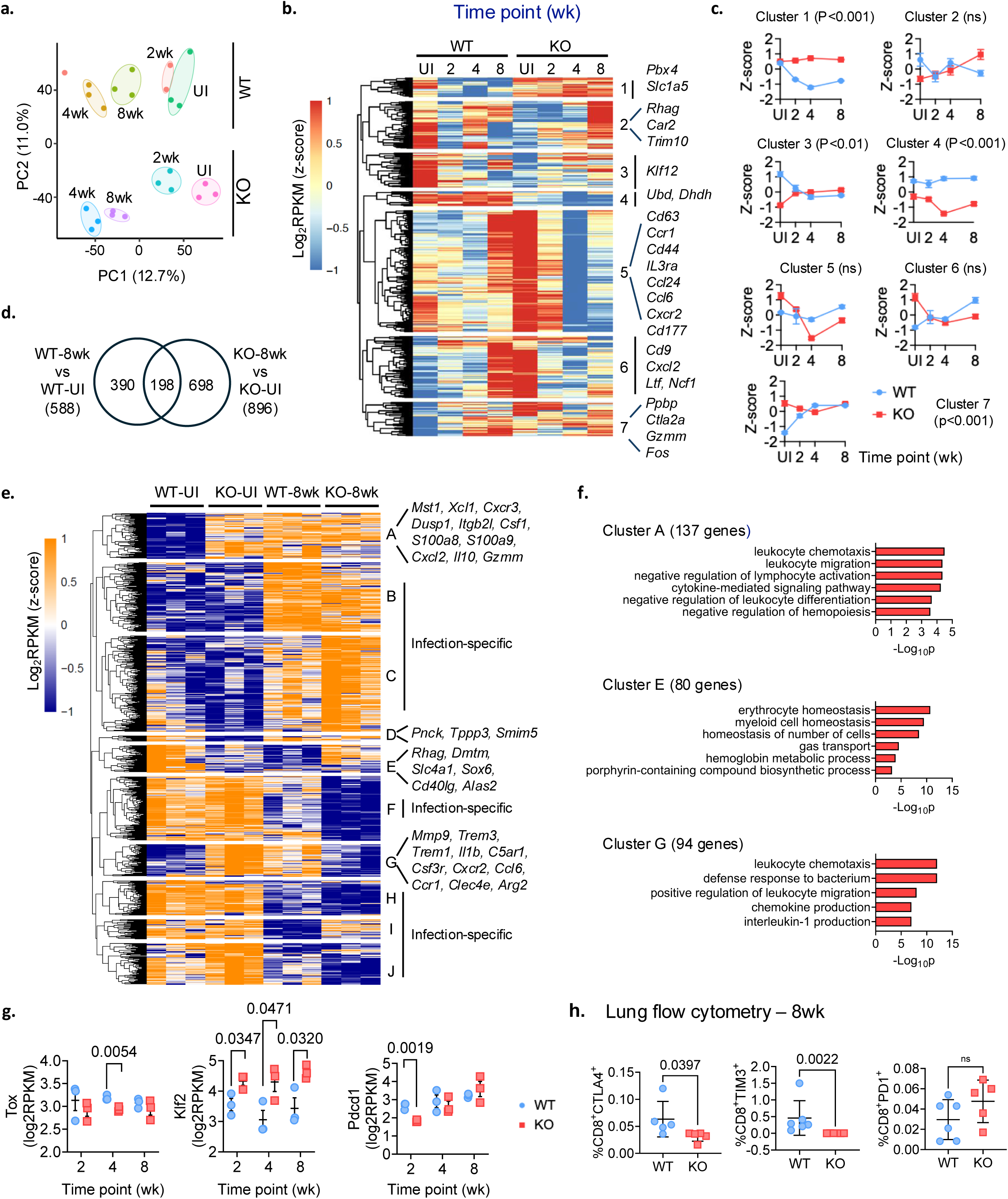
Transcriptomic profiling of WT and *Nr4a1*^⁻/⁻^ (KO) CD8^+^ T cells identify distinct gene expression programs. **a**, PCA plot of bulk RNA-seq libraries from uninfected (UI) and infected (2, 4, 8 wk) spleen CD8^+^ T cells. Samples from the same group are circled manually except for an outlier in WT group at 2 wk time point. **b**, Heatmap of time-course DEGs (*n* = 463) between *Nr4a1^-/-^* and WT CD8^+^ T cells. Union of *Nr4a1^-/-^* vs WT DEGs identified at 2, 4, 8wk using UI mice as baseline was used. Color bar denotes log_2_ RPKM values averaged for each group and z-score normalized. **c**, Mean z-scores (± s.e.m) for each cluster in panel b across infection time points. Trends in cluster 1, 4, and 7 are significant (*P*<0.05, two-way ANOVA with Šídák’s multiple-comparisons test). **d**, Venn diagram showing DEGs overlap between 8 wk and UI spleens in WT and KO mice. **e**, Heatmap of the union of 8wk vs UI DEGs (*n* = 1,286) from panel d, showing ten clusters (A-J) derived by k-means. Color bar denotes log_2_ RPKM values that were z-score normalized. **f**, GO biological processes enriched in clusters A, E and G. **g**, Expression kinetics of exhaustion-related genes *Tox*, *Klf2* and *Pdcd1* in infected WT and KO lungs. *P values* by unpaired two-tailed *t*-test. **h**, Flow-cytometric analysis of CD8^+^ T cells at 8 wk in WT and KO mice (*n* = 5); *P values*, two-tailed Mann-Whitney test.

To compare genotypes at specific time points, we analyzed DEGs relative to uninfected controls. At 4 wk, WT CD8^+^ T cells exhibited 782 DEGs (WT-4wk vs WT-UI), whereas *Nr4a1*^⁻/⁻^ CD8^+^ T cells showed 1330 DEGs (KO-4wk vs KO-UI, Extended Data Fig. 2a), with 337 shared. Heatmap visualization identified ten clusters (cluster i-x, Supplementary Table 2), four of which (clusters ii, vi, viii and ix) were infection-specific (Extended Data Fig. 2b). Only clusters i and v showed enrichment of gene ontology (GO) (Extended Data Fig. 2c). Cluster i genes (n = 147) were elevated in *Nr4a1*^⁻/⁻^-UI compared to WT-UI, and encoded for T-helper 1 cytokine production (*Rora*, *Tbx21*, *Xcl1*, *Klrc*1, *Il18rap*) and cytokines/chemokines pathways (*Ccl5*, *Ccl4*, *Xcr1*, *Cxcr3*). The cluster i genes were also upregulated in both infected WT and *Nr4a1*^⁻/⁻^ CD8^+^ T cells (i.e KO-4wk and WT-4wk, Extended Data Fig. 2b). In contrast, cluster v genes (n = 229) showed upregulation only in *Nr4a1*^⁻/⁻^ UI (KO-UI) group, and encoded proteins for leukocyte migration (*Anxa1*, *Cd177*, *Gdf15*, *Ccl6*, *Cxcr1*, *Cxcl13*) and adhesion (*Sirpb1a*, *Sirpb1b*, *Lilrb4a*, *Lilrb4b*, *Ptafr*) (Extended Data Fig. 2c).

At 8 wk after infection, the DEG patterns were similar (Fig. 2d-f and Supplementary Table 2). Greater number of DEGs were identified in *Nr4a1^-/-^*-CD8^+^ T cells (896 vs 588; Fig. 2d). Unsupervised clustering identified ten gene clusters (cluster A to J, Supplementary Table 2); six of which (cluster B, C, F, H, I, J) were infection-specific (Fig. 2e). Clusters A, E and G showed enrichment of GO biological processes (Fig. 2f). Cluster A genes (higher expression in *Nr4a1^-/-^* UI) encoded proteins for leukocyte chemotaxis and migration (Fig. 2e,f). However, Cluster E and G genes (differentially expressed in *Nr4a1^-/-^*-CD8^+^ T cells vs WT), encoded proteins for leukocyte homeostasis, differentiation, chemotaxis and activation (*Cd40lg*, *Alas2*, *Ccl6*, *Ccr1*, *Arg2*, and *Sox6*)^27, 28, 29, 30^.

Given that NR4A1 regulates transcription factors such as *Klf2*, *Irf4*, and *Tox* to limit T cell activation and promote exhaustion ^13, 17, 18, 31, 32^, we examined these genes. *Nr4a1*^⁻/⁻^ CD8^+^ T cells displayed reduced *Tox* and increased *Klf2* (Fig. 2g) accompanied by decreased *Pdcd1* (Fig. 2g). Consistently, we found lower percentages of CTLA4^+^ and TIM3^+^ exhausted CD8^+^ T cells in *Mtb*-infected *Nr4a1*^⁻/⁻^ lungs at 8wk (Fig. 2h). Together, these analyses show that NR4A1 deficiency promotes an activated and less exhausted CD8^+^ T cells during *Mtb* infection.

### NR4A1 deficiency augments CD8^+^ T cell driven protection in TB

Based on the gene expression data, we questioned whether *Nr4a1^⁻^*^/⁻^ CD8^+^ T cells confer superior protection against *Mtb* infection in vivo. To test this, splenic CD8^+^ T cells from WT and *Nr4a1^⁻^*^/⁻^ mice were adoptively transferred into *Rag1*^⁻/⁻^ mice that were infected one day later with *Mtb* Erdman (Fig. 3a). Comparison was made with transferred WT and *Nr4a1^⁻^*^/⁻^ CD4^+^ T cells (Fig. 3a). Transferred CD4^+^ T cells from either genotype affected bacterial load to a similar extent (Fig. 3b). In contrast, *Rag1*^⁻/⁻^ recipients of *Nr4a1*^⁻/⁻^ CD8^+^ T cells exhibited significantly lower pulmonary bacterial burden compared to those receiving WT CD8^+^ T cells, with a reduction of CFU comparable to transferred CD4^+^ T cells (Fig. 3b,c). Histological analysis revealed that *Rag1*^⁻/⁻^ mice receiving *Nr4a1*^⁻/⁻^ CD8^+^ T cells had smaller and less consolidated lesions compared to mice receiving either WT CD8^+^ T cells or no-transfer controls (Fig. 3c, compilation graph). *Rag1*^⁻/⁻^ mice receiving *Nr4a1*^⁻/⁻^ CD4^+^ T cells also had smaller lesions similar to WT CD4^+^ T cells (Fig. 3c). Cytokine analysis of lung homogenates revealed a significantly different inflammatory profile in *Rag1*^⁻/⁻^ mice that received *Nr4a1*^⁻/⁻^ CD8^+^ T cells compared with those receiving WT CD8^+^ T cells (CD8-KO vs CD8-WT, Fig. 3d). Importantly, *Rag1*^⁻/⁻^ mice receiving *Nr4a1*^⁻/⁻^ CD8^+^ T cells demonstrated increased CCL3 (Fig 3d), which regulate CD8^+^ T cell migration and effector functions^33, 34^. CD8^+^ T cells are primarily known for their antigen-specific cytotoxicity, but they can also exhibit an antigen-nonspecific antimicrobial activity^35^. We evaluated this by testing the anti-*Mtb* activity of *Nr4a1*^⁻/⁻^ CD8^+^ T cells in vitro. *Mtb* (H37Rv-lux or CDC1551) infected thioglycolate-elicited peritoneal macrophages (TEPMs) were co-cultured with CD8^+^ T cells from either BCG-vaccinated or unvaccinated WT and *Nr4a1*^⁻/⁻^ mice (Fig. 3e). CD8^+^ T cells from both unvaccinated and BCG-vaccinated *Nr4a1*^⁻/⁻^ mice showed significantly greater suppression of *Mtb* replication compared to their WT counterparts (Fig. 3f,g). Notably, CD8^+^ T cells from BCG-vaccinated *Nr4a1*^⁻/⁻^ mice showed better protection compared with unvaccinated *Nr4a1*^⁻/⁻^ mice. These results demonstrated that NR4A1-deficient CD8^+^ T cells restrict *Mtb* in both antigen-specific and antigen-nonspecific manners, indicating a CD8^+^ T cell–intrinsic role for NR4A1 in limiting bacterial control.

**Fig. 3.**
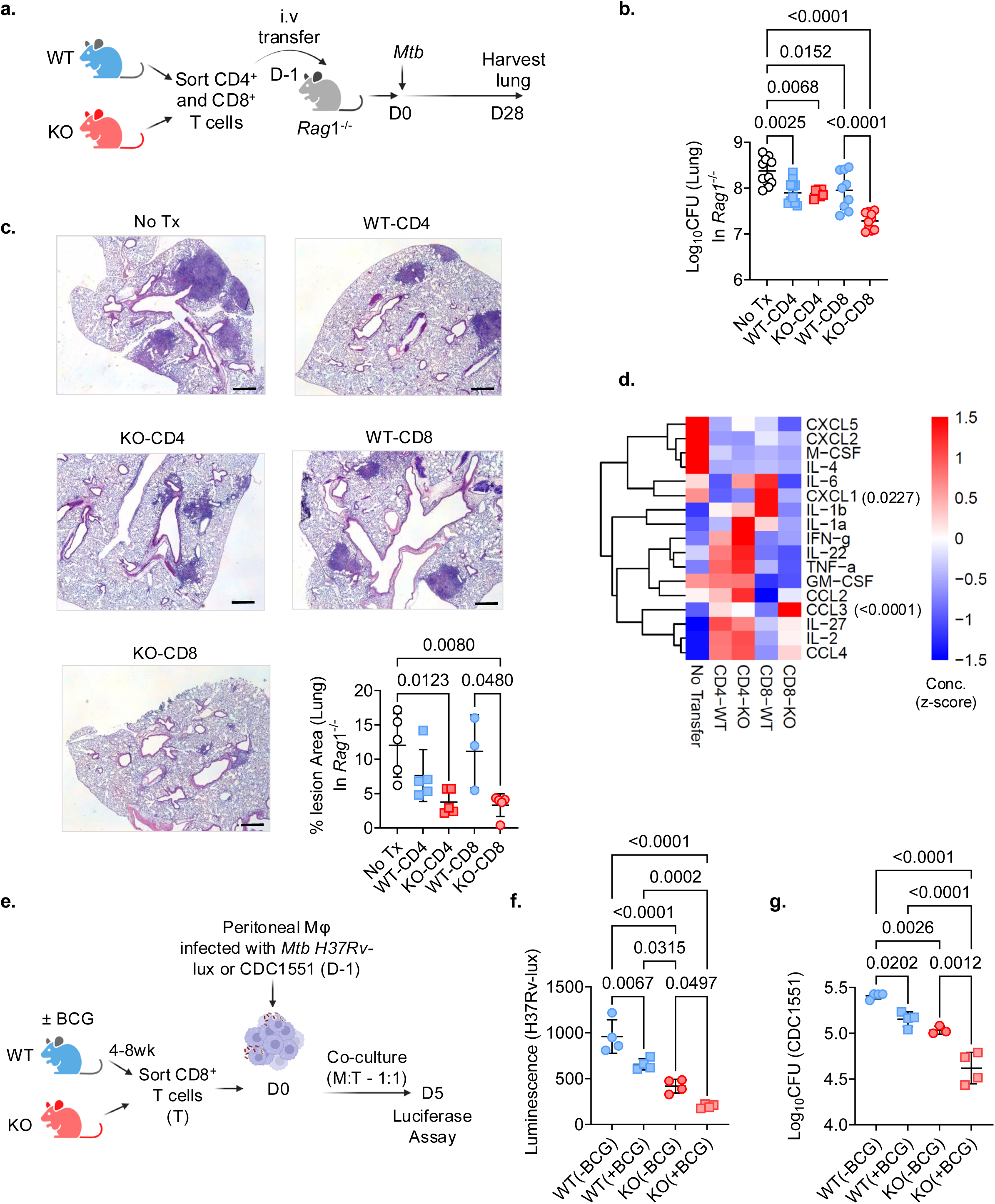
NR4A1 regulates the function of CD8^+^ T cells in *Mtb-*infected mice. **a**, Schematic of the adoptive transfer experiment—Magnetically sorted CD4^+^ and CD8^+^ T cells isolated from naïve WT and *Nr4a1*^−/−^ mice were transferred i.v. into *Rag1*^-/-^ recipients. The following day, mice were infected with *Mtb* Erdman and then harvested on day 28. **b**, Lung bacterial load in *Rag1*^-/-^ mice. No Tx - *Rag1*^-/-^ mice without transferred cells; WT-CD4 and KO-CD4 - *Rag1*^-/-^ mice receiving CD4^+^ T cells from WT and KO mice respectively; WT-CD8 and KO-CD8 - *Rag1*^-/-^ mice receiving CD8^+^ T cells from WT and KO mice respectively. Data pooled from 2 independent experiments (*n* = 5 per group). **c**, Representative H&E-stained lung sections from the *Rag1*^-/-^ recipients as in b. Bottom right: quantification of TB lesion area as a percentage of total lung area (*P* = 0.0480; one-way ANOVA). **d**, Cytokine and chemokine analysis in lung homogenates from *Rag1*^-/-^ mice in B. **e**, Schematic of the in vitro co-culture experiment. WT or *Nr4a1*^-/-^ mice were vaccinated with BCG or sham vaccinated with sterile saline. After 4–8 wk, CD8^+^ T cells were sorted and co-cultured for 5 days in a 1:1 ratio with WT peritoneal macrophages that were infected with *Mtb* H37Rv-lux or CDC1551 strain (MOI = 1). **f**, Bacterial growth expressed as relative luminescence units (RLU) in co-cultures of infected macrophages with CD8^+^ T cells from vaccinated or unvaccinated (± BCG) WT and *Nr4a1*^−/−^ mice. **g**, Same as f, but bacterial burden measured as CFU of CDC1551. In f and g each dot represents data from one mouse; *P values* in b, d, f and g, one-way ANOVA. Data in a-c and e-g are representative of 2 and 3 independent experiments, respectively.

### scRNA-seq reveals reprogramming of *Nr4a1⁻^/⁻^* lung CD8^+^ T cells towards enhanced effector and cytotoxic states

To understand how NR4A1 influences CD8^+^ T cell responses in the lung during *Mtb* infection, we performed single-cell RNA sequencing (scRNA-seq) on lung T cells from WT and *Nr4a1*^⁻/⁻^ mice at 4 wk (Fig. 4a). CD8^+^ T cells from the lung of uninfected mice served as controls. Based on the expression of 30 genes and unsupervised clustering, Uniform Manifold Approximation and Projection (UMAP) analysis of 10,452 lung CD8^+^ T cells identified 16 CD8^+^ T cell clusters/subsets, including naïve, effector (Teff), effector memory (Tem), memory (Tm), tissue-resident memory (Trm), exhausted (Tex), and proliferating cells (Tprolif) (Fig. 4b and Extended Data Fig. 3a). Compared to *Mtb*-infected WT mice, *Mtb*-infected *Nr4a1*^⁻/⁻^ mice exhibited increased frequency of cytotoxic effector subsets, particularly *Gzmk*^hi^ Tem (cluster g) and *Gzma*^hi^ Teff (cluster m) populations (Fig. 4c,d). In contrast, CD8^+^ T cell subsets *Il2ra*^hi^*Xcl1*^+^ Treg, *Furin*^hi^ Trm, and *Ifitm1*^+^ Trm were decreased in *Mtb*-infected *Nr4a1*^⁻/⁻^ mice (Extended Data Fig. 3b). DEG analysis revealed strong upregulation of cytotoxicity-associated genes such as *Gzma*, *Gzmb*, *Nkg7*, *Klrk1*, and *Ccl5* in *Gzmk*^hi^ Tem and *Gzma*^hi^ Teff subsets of *Nr4a1*^⁻/⁻^ CD8^+^ T cells (Fig. 4e, Supplementary Table 3). We noted that *Gzma* and *Nkg7* were upregulated in total *Nr4a1*^⁻/⁻^ CD8^+^ T cells both uninfected and *Mtb*-infected conditions relative to WT counterpart (pseudobulk gene expression, Fig. 4f,g). GO analysis revealed significant enrichment of multiple immune effector-related biological processes in *Gzmk*^hi^ Tem and *Gzma*^hi^ Teff *Nr4a1*^⁻/⁻^ CD8^+^ T cells (Fig. 4h). The top enriched pathways among upregulated DEGs were cell killing and natural killer (NK) cell-mediated cytotoxicity, whereas among downregulated DEGs was cellular response to interferon-beta. Notably, suppression of type 1 IFN signaling has been associated with TB protection^36^. Chemotaxis-related pathways were also enriched, suggesting that NR4A1-deficient CD8^+^ T cells possess altered migratory capacity, consistent with the bulk RNA-seq analysis (Fig. 2). Collectively, scRNA-seq indicated that NR4A1-deficient CD8^+^ T cells express an altered transcriptional program characterized by enhanced (i) effector function, (ii) innate-like activation, and (iii) potential for improved trafficking.

**Fig. 4.**
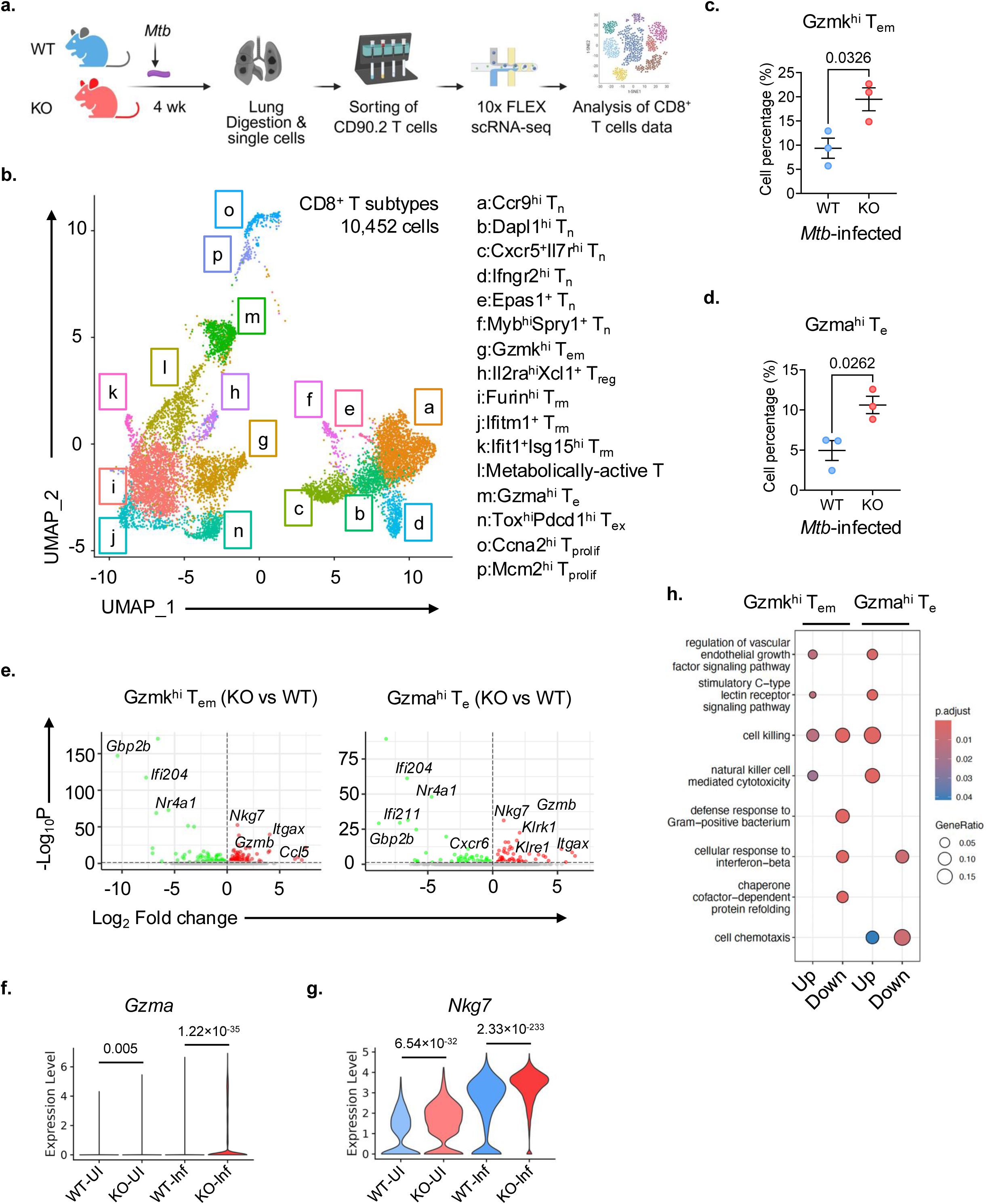
NR4A1 deficient CD8^+^ T cells display enriched expression of cytotoxicity-associated genes. **a**, Schematic of scRNA-seq workflow for analyzing lung CD8^+^ T cell dataset in UI and infected WT and *Nr4a1*^−/−^ (KO) mice. **b**, UMAP visualization of CD8^+^ T cell subsets from infected lungs: T_n_ (naïve), T_em_ (effector memory), T_m_ (memory), T_rm_ (tissue-resident memory), T_e_ (effector), T_ex_ (exhausted), and T_prolif_ (proliferating). Cluster identities are annotated. **c,d**, Frequencies of *Gzmk*^hi^ T_em_ (cluster g) and *Gzma*^hi^ T_e_ (cluster m) cells in infected WT and KO mice. *P* values, unpaired *t*-test. **e**, Volcano plots showing DEGs in *Gzmk*^hi^ T_em_ (left) and *Gzma*^hi^ T_e_ (right) comparing *Nr4a1*^−/−^ versus WT cells. Genes upregulated in CD8^+^ T cells from KO mice are shown in red; downregulated genes in green. **f,g**, Violin plots depicting pseudogene expression of *Gzma* and *Nkg7* among CD8^+^ T cells from uninfected WT (WT-UI), uninfected KO (KO-UI) and their infected counterpart (WT-Inf and KO-Inf). **h**, Enrichment analysis plots displaying enhanced pathways among DEGs of *Nr4a1*^−/−^ versus WT identified in *Gzmk*^hi^ T_em_ and *Gzma*^hi^ T_e_ clusters (dot sizes reflect the gene ratio in the pathway).

### NR4A1 negatively regulates NKG7 expression and limits CD8^+^ T cell–mediated cytotoxicity during *Mtb* infection

To identify the mechanisms underlying the enhanced cytolytic activity of *Nr4a1*^⁻/⁻^ CD8^+^ T cells, we examined the common DEGs across all 16 CD8^+^ T cell clusters in the scRNA-seq dataset (Fig. 4b). This identified *Nkg7* and *Gvin1* as the top two upregulated genes shared by 12 of the 16 CD8^+^ T cell clusters (Fig. 5a). The longitudinal bulk RNA-seq analysis from *Mtb*-infected mice (Fig. 2) also confirmed elevated *Nkg7* mRNA expression in splenic *Nr4a1*^⁻/⁻^ compared to WT CD8^+^ T cells (Extended Data Fig. 4a). This suggested that *Nkg7* expression might be regulated by NR4A1. NR4A1 regulates target gene expression by binding to NGFI-B-response elements within promoter regions^37^. Analysis of the *Nkg7* promoter using the JASPAR database predicted two NR4A1-binding motifs (AAAGGTCA) (Fig. 5b). ChIP-qPCR analysis on splenic CD8^+^ T cells from WT mice confirmed NR4A1 binding at both predicted sites (Fig. 5c). *Nkg7* encodes a granule associated effector protein essential for T cell cytotoxicity that plays a crucial role in the formation, trafficking and release of lytic granules^38, 39^. Having established that NR4A1 binds and represses *Nkg7* transcription, we therefore examined whether this regulation influences the NKG7 expression in *Nr4a1*^⁻/⁻^ CD8^+^ T cells. Interestingly, flow-cytometric analysis demonstrated higher percentage of CD8^+^NKG7^+^, CD8^+^Perforin^+^ and CD8^+^CD107a^+^ T cells in the lungs of *Nr4a1*^⁻/⁻^ mice compared to WT mice (Fig. 5d-f), supporting an enrichment of the cytolytic program among CD8^+^ T cells in the absence of NR4A1.

**Fig. 5.**
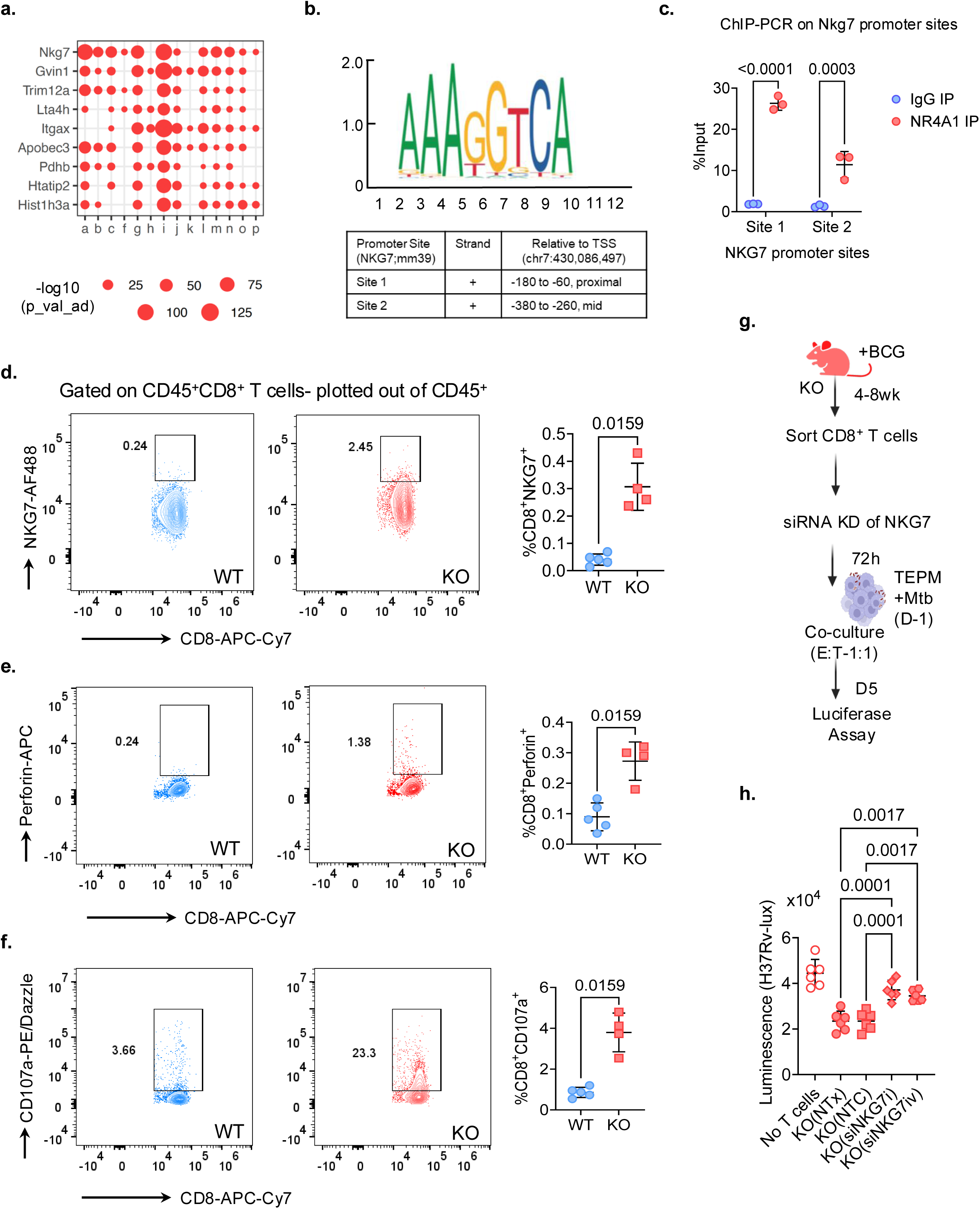
NKG7 mediates the enhanced cytotoxic and protective function of *Nr4a1*^⁻/⁻^ CD8^+^ T cells. **a**, Dot plot showing representative DEGs associated with cytotoxic effector function (*Nkg7, Gzmb, Prf1, Itga2,* and others) across 14 CD8^+^ T cell clusters identified from scRNA-seq data. Circle size represents the fraction of cells expressing each gene and color intensity reflects the average expression level. **b**, Two genomic sites, relative to the transcription start site (TSS), within the *Nkg7* promoter predicted to have NR4A1-binding motif (AAAGGTCA), using JASPAR. **c**, ChIP–qPCR validation of NR4A1 binding on *Nkg7* promoter in WT splenic CD8^+^ T cells. Results are expressed as % input DNA. Data presented as mean ± s.d.; *n* = 3 mice per group; P values bytwo-way ANOVA. **d-f**, Representative flow-cytometry plots showing expression of NKG7 (d), Perforin (e) and CD107a (f) in CD8^+^ T cells from WT and *Nr4a1*^⁻/⁻^ mice. Respective quantification of cells among CD8^+^ T cells is indicated on right side. Data presented as mean ± s.d.; *n* = 4 mice per group; *P* values - two-tailed Mann–Whitney test. **g**, Schematic outlining the experimental design for siRNA-mediated knockdown of *Nkg7* in CD8^+^ T cells isolated from BCG vaccinated *Nr4a1*^−/−^ mice. CD8^+^ T cells were either left untreated (NTx), treated with non-targeting control (NTC), or transfected with *Nkg7*-specific siRNAs (siRNAi and siRNAiv), followed by 5-day co-culture with *Mtb* H37Rv-lux infected TEPMs. **h**, Luminescence readout of bacterial burden (H37Rv-lux) in co-cultures with the indicated CD8^+^ T cell treatments. Data presented as mean ± s.d; Each dot represents an individual mouse; *P value* one-way ANOVA. Data (d-h) is from 2-3 independent experiments.

Next, we tested the functional relevance of increased NKG7 in *Nr4a1^-/-^* CD8^+^ T cells using siRNA knockdown in an in vitro *Mtb* co-culture system. CD8^+^ T cells isolated from BCG-vaccinated *Nr4a1^-/-^*mice were transfected with *Nkg7*-specific siRNAs (siNKG7i and siNKG7iv), a non-targeting control (NTC) siRNA, or mock-transfected, and subsequently co-cultured with H37Rv-lux-infected TEPMs for five days (Fig. 5g and Extended Data Fig. 4b). Knockdown of *Nkg7* abolished the ability of *Nr4a1^-/-^* CD8^+^ T cells to suppress *Mtb* growth (Fig. 5h), indicating that NKG7 is a critical effector of bacterial control by *Nr4a1^-/-^*CD8^+^ T cells. Together, these findings defined an NR4A1–NKG7 axis governing CD8^+^ T cell cytotoxicity and anti-*Mtb* defense.

### *Nr4a1*^⁻/⁻^ mice show increased infiltration of the CD8^+^ T cells into TB lesions

We next investigated the effect of NR4A1 ablation on CD8^+^ T cell localization in lung TB lesions. Immunofluorescence (IF) microscopy and the GeoMx Digital Spatial Profiler (DSP) whole transcriptome atlas (WTA) assay^40^ were used to map the spatial localization of T cells in lung sections from WT and *Nr4a1*^⁻/⁻^ mice at 4 wk (Fig. 6a). IF staining for CD8^+^ and CD4^+^ T cells revealed distinct spatial distribution patterns of T cell subsets within TB lesions (Fig. 6b). In WT mice, CD8^+^ cells were largely restricted to the lesion periphery, while in *Nr4a1*^⁻/⁻^ mice, CD8^+^ cells were more prominently localized within the lesion core (Fig. 6b). In contrast, CD4^+^ T cell localization was unchanged between the two genotypes. Quantitative analysis of IF images confirmed a significantly higher density of CD8^+^ T cells within the lesion core of *Nr4a1*^⁻/⁻^ mice compared to WT controls (Fig. 6c). In contrast, the number of CD4^+^ cells in the same region were comparable between the two groups (Fig. 6d). These observations, together with our flow cytometry data (Fig. 1h,i), suggested that NR4A1 shapes lesion architecture by limiting CD8^+^ T cell infiltration into the lesion core.

**Fig. 6.**
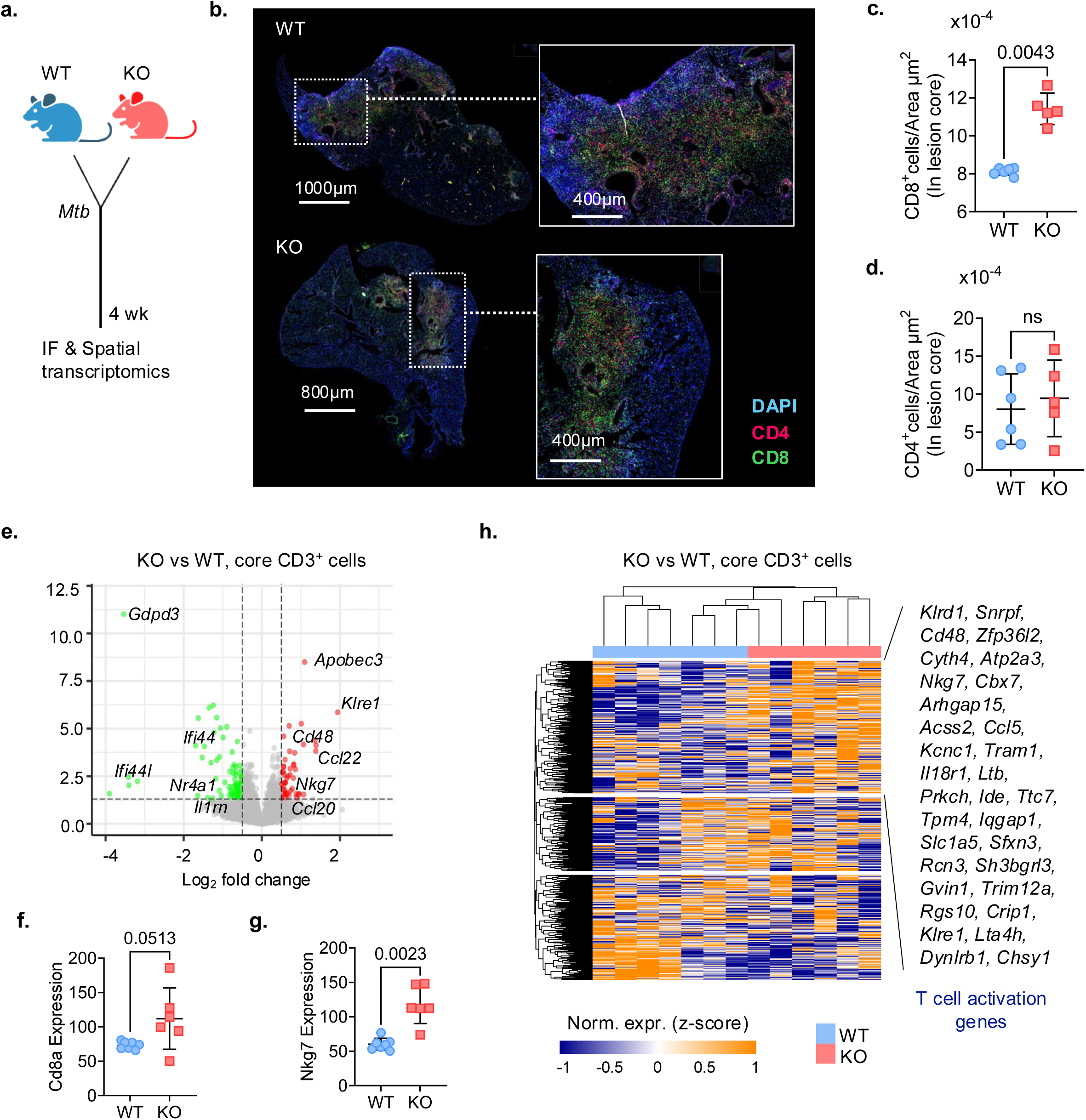
*NR4A1*^−/−^ mice have increased infiltration of the CD8^+^ T cells in the lesion core. **a**, Schematic - WT and *Nr4a1*^−/−^ (KO) mice were infected with low-dose *Mtb* Erdman and euthanized at 4 wk; lungs were harvested for immunofluorescence (IF) and spatial transcriptomics (see Methods). **b**, Representative IF images of lung sections; zoom panels show areas of lesion cores infiltrated by CD8^+^ (green) and CD4^+^ (red) T cells in the WT (top) and KO (bottom) mice. DAPI is coded in blue. **c,d**, Plots showing number of CD8^+^ cells or CD4^+^ cells per µm² in lesion cores of *Mtb*-infected *Nr4a1*^−/−^ versus WT mice. *P* values, two-tailed Mann–Whitney test, two independent experiments. **e**, Volcano plot showing DEGs in spatially resolved CD3^+^ T cells isolated from lungs of *Mtb*-infected KO versus WT mice (See Extended Data Fig. 5). Genes significantly upregulated in KO T cells (adjusted *P* < 0.05, log_2_FC > 1) are shown in red; downregulated genes (log_2_FC < −1) in green; non-significant genes in gray. Notable DEGs are annotated, including increased expression of cytotoxicity-related genes such as *Nkg7*. **f,g**, Mapping of *Cd8a* (f) and *Nkg7* (g) mRNA expression in CD3^+^ T cell–enriched regions from lungs of WT and KO mice. Each point represents normalized *Cd8a* or *Nkg7* transcript abundance from an individual ROI. *P value* two-tailed Mann–Whitney test. **h**, Heatmap of the pseudobulk DEGs identified from total lung CD8^+^ T cell scRNA-seq (Fig. 5b), evaluated in core CD3^+^ ROIs from GeoMx spatial transcriptomics of *Mtb*-infected lungs. Representative cytotoxic and activation genes enriched in KO—*Klrd1, Cd48, Nkg7, Ccl5, Klre1*—are highlighted on right side. See Methods for DEG definition, normalization and clustering parameters.

To uncover the underlying transcriptional programs associated with the intralesional CD8^+^ T cell shift, we performed spatial transcriptomic profiling on CD3^+^ T cells in the core and periphery of lung lesions from *Mtb*-infected WT and *Nr4a1*^⁻/⁻^ mice (Extended Data Fig. 5a,b). We confirmed the presence of acid-fast bacilli (AFB) bacilli in the core of lung lesions on an adjacent serial section (Extended Data Fig. 5c). The AFB region and CD3^+^ staining guided the region of interest (ROI) selection from the lesion core and periphery (Extended Data Fig. 5a-c). DEG analysis among CD3^+^ T cells from core ROIs revealed upregulation of effector and cytotoxicity-related genes (*Ccl20*, *Ccl5*, *Cd48*, *Klre1*, and *Klrd1*) in *Nr4a1*^⁻/⁻^ ROIs compared to WT ROIs (Fig. 6e, Supplementary Table 4). Core CD3^+^ T cells from *Nr4a1*^⁻/⁻^ mice also exhibited higher expression of *Cd8a* and *Nkg7* (Fig. 6f,g), consistent with enhanced CD8^+^ T cell abundance and their cytotoxic activation in *Nr4a1*^⁻/⁻^ mice (Fig. 1, 4, 5). In contrast, *Cd4* expression levels in core CD3^+^ T cells remained unaltered (Extended Data Fig. 4d). Unsupervised hierarchical clustering of core CD3^+^ T cell–associated ROIs using pseudobulk CD8^+^ T cell DEGs from scRNA-seq data clearly separated WT from *Nr4a1*^−/−^ mice, (Fig. 6h and Extended Data Fig. 5e). This clustering further demonstrated high expression of genes related to T cell activation, such as *Cd48*, *Ccl5*, *Klrd1*, in *Nr4a1*^⁻/⁻^ mice. Peripheral CD3^+^ T cell ROIs profiled by GeoMx DSP also showed elevated expression of *Cd8a, Cd4* and *Nkg7* in *Nr4a1*^⁻/⁻^ mice relative to WT counterparts, but they could not clearly segregate WT from *Nr4a1*^−/−^ mice (Extended Data Fig. 5f,g). Collectively, these findings revealed that NR4A1 acts as a transcriptional brake on both the spatial infiltration and cytotoxic programming of CD8^+^ T cells within pulmonary TB lesions.

### Lung *NR4A1*^lo^ CD8^+^ T cells from human and macaque display activated and cytotoxic gene signatures in TB

To establish that NR4A1 expression indeed stratifies functional states of CD8^+^ T cells in TB, we reanalyzed published scRNA-seq datasets from the lungs of *Mtb*-infected cynomolgus macaques (macaques) and active TB patients^41, 42^. CD8^+^ T cells from TB granulomas of infected macaques were categorized into four subsets, T_EMRA_, Tc17, Effector (T_Eff_) and *Gzmk*^i^ (Extended Data Fig. 6a); with Tc17 cells showing the highest expression of *Nr4a1* mRNA (Fig. 7a and Extended Data Fig. 6b). Comparing gene expression of macaque *Nr4a1*^lo^ and *Nr4a1*^hi^ CD8^+^ T cells revealed upregulation of cytotoxic and activation-associated genes, including *Nkg7*, *Gzmb*, *Gzmh*, *Gzma*, *Gnly*, *Ccl5*, *Prf1*, in the *Nr4a1*^lo^ subset (Fig. 7b, Supplementary Table 5). GO analysis of genes upregulated in *Nr4a1*^lo^ cells showed an enrichment for interleukin-2 signaling, T cell receptor signaling, CTL-mediated killing, and antigen processing/cross-presentation biological processes (Fig. 7c). A similar gene expression pattern reflecting an activated and cytotoxic phenotype of lung *Nr4a1*^lo^ CD8^+^ T cells was observed in humans with active TB (Fig. 7d,e, Extended Data Fig. 6c,d). Consistent with the macaque data, human lung *Nr4a1*^lo^ CD8^+^ T cells displayed increased expression of effector and cytotoxic genes such as *Nkg7*, *Gzmb*, *Gzmk*, *Prf1*, Ccl5, which was highlighted in GO enrichment analysis (Fig. 7e,f, Supplementary Table 5). These cross-species analyses validated the relevance of mouse data and supported the hypothesis that NR4A1 is transcriptional checkpoint whose expression inversely correlates with cytotoxic effector programming.

**Fig. 7.**
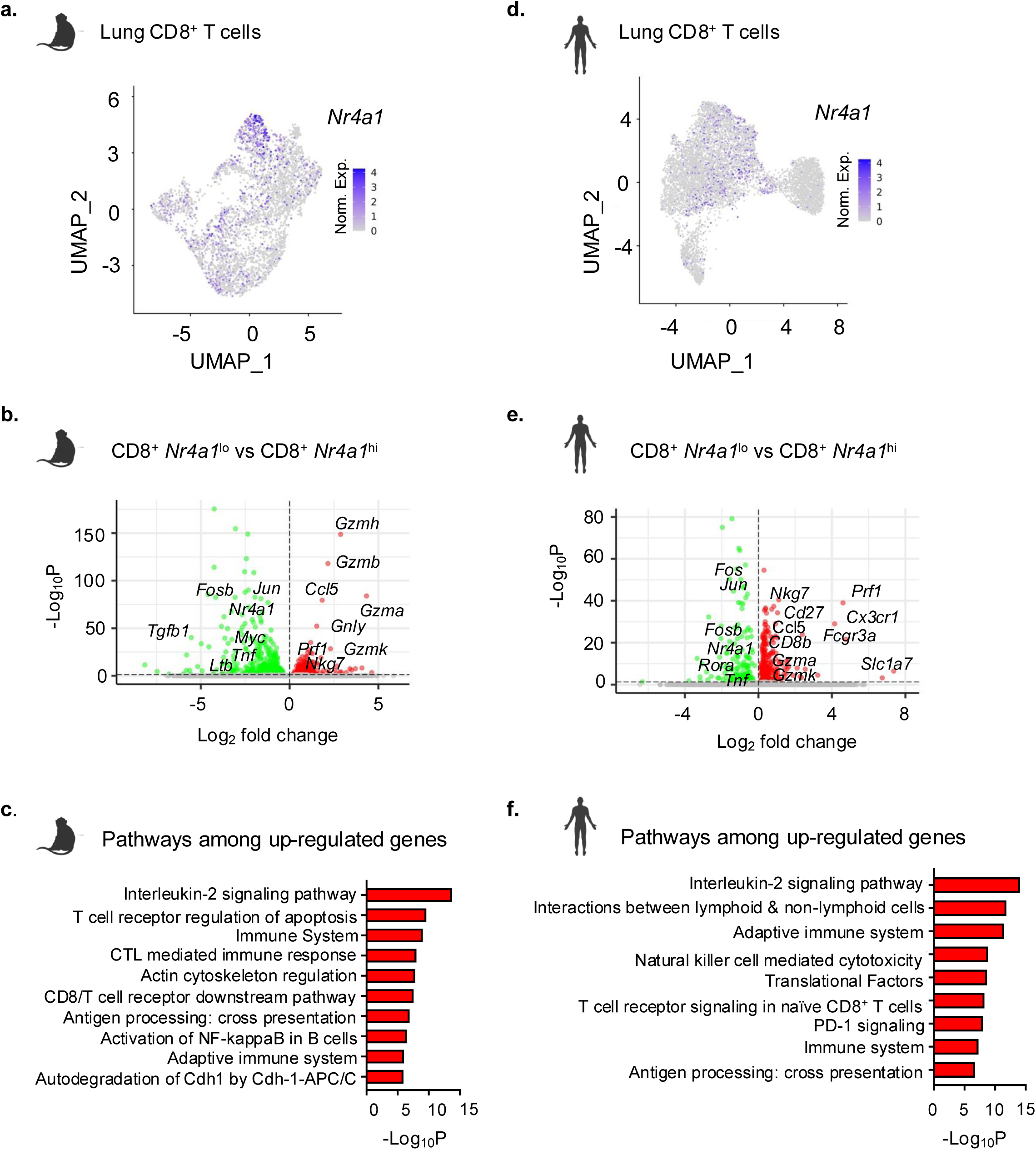
Lung *Nr4a1*^lo^ CD8^+^ T cells from *Mtb* infected macaque and human display activated and cytotoxic transcriptional program. Re-analysis of single-cell RNA-seq data from *Mtb-*infected cynomolgus macaque ^41^ and active TB patient ^42^ lungs identified CD8^+^ T cells expressing low (*Nr4a1*^lo^) and high (*Nr4a1*^lo^) *Nr4a1* gene. **a**, Feature plot showing *Nr4a1* expression across CD8^+^ T cell clusters in macaque lung. **b**, Volcano plot showing DEGs in macaque *Nr4a1^lo^*versus *NR4A1^hi^* CD8^+^ T cells, revealing up-regulation of cytotoxic genes (*Nkg7*, *Gzmb*, *Gzmh*, *Gnly*, *Prf1*, *Ccl5*). **c**, Pathway enrichment (using Bioplanet) among up-regulated genes in b. **d**, Feature plot depicting *Nr4a1* expression across CD8^+^ T cell clusters in the lung of active TB patient. **e**, Volcano plot showing DEGs in TB patient lung *Nr4a1^lo^* versus *NR4A1^hi^* CD8^+^ T cells. **f**, Pathway enrichment analysis among up-regulated genes in e.

### NR4A1 inhibition improves TB control in mice

We next asked whether pharmacological inhibition of NR4A1 in WT mice could recapitulate the TB protective phenotype observed in *Nr4a1*^−/−^ mice (Fig. 1). To test this, *Mtb*-infected C57BL/6 mice were treated with the NR4A1 antagonist DIM-C-pPhCO₂Me (DIM-C; 40 mg/kg/day)^43, 44^ starting on day 7 after infection (Fig. 8a). At the experimental endpoint (day 21), DIM-C–treated mice exhibited significantly lower pulmonary bacterial loads compared to vehicle-treated controls (Fig. 8b). Moreover, H&E staining revealed that DIM-C treatment remodeled lung pathology, yielding smaller and more diffuse lesions, in contrast to the large, well-demarcated lesions in the vehicle-treated mice (Fig. 8c). Quantitative analysis confirmed a significant reduction in lesion area in DIM-C–treated mice (Fig. 8d). DIM-C treatment significantly reduced *Nr4a1* transcript levels while concomitantly upregulating *Nkg7* expression in the lung tissue (Fig. 8e,f). Importantly, IF analysis demonstrated that DIM-C treatment increased the infiltration of CD8^+^ T cells into the core of lung lesions (Fig. 8g,h), mirroring the spatial redistribution of *Nr4a1*^⁻/⁻^ CD8^+^ T cells (Fig. 6b,c). In contrast, CD4^+^ T cell distribution was comparable between groups (Fig. 8g,i). Taken together, our findings demonstrate that inhibition of NR4A1 drives the redistribution of activated CD8^+^ T cells with enhanced cytotoxic potential in the lung during TB (Fig. 8j).

**Fig. 8.**
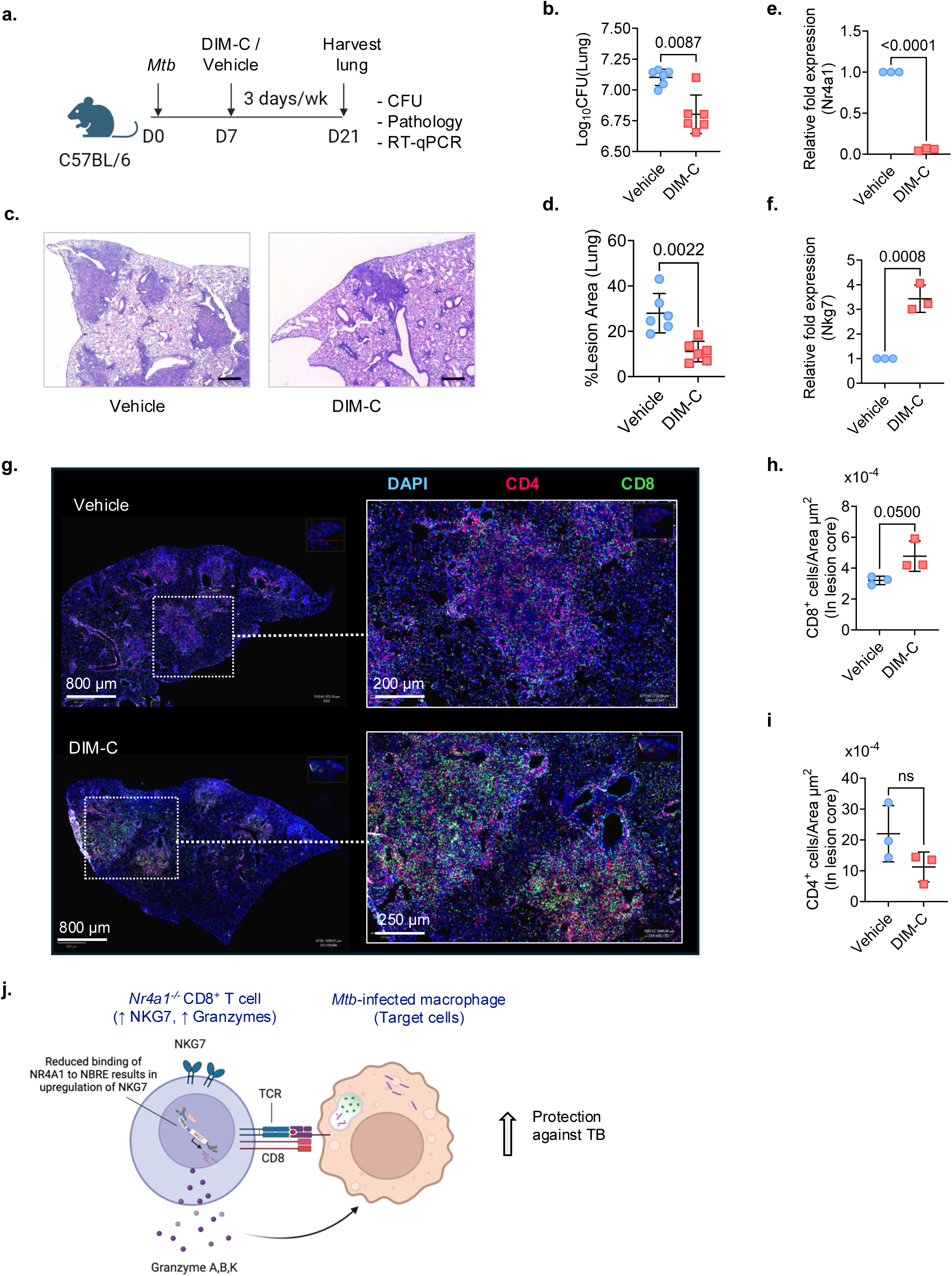
Pharmacological inhibition of NR4A1 enhances CD8^+^ T cell infiltration and protection during *Mtb* infection. **a**, Experimental schematic: C57BL/6 mice (6–8 weeks old) were administered the NR4A1 antagonist DIM-C-pPhCO2Me (DIM-C; 40 mg/kg/day) via oral gavage every other day for 21 days following low-dose aerosol infection with *Mtb* Erdman. Control mice received vehicle (corn oil) alone. **b**, Lung bacterial burden in DIM-C treated (red) versus vehicle treated (blue) mice. Data are mean ± s.d. *P values*, two-tailed Mann-Whitney test. **c**, Representative lung histology images (H&E) in DIM-C treated mice (right) versus control (left) (scale bar = 100 µm). **d**, Quantification of granulomatous lesion area (% lung area) from C*. P* value, two-tailed Mann–Whitney test. **e,f**, RT-qPCR of *Nr4a1* (e) and *Nkg7* (f) transcript levels in the lung normalized to β-actin. **g**, Representative IF images of lung sections stained for CD4 (red), CD8 (green), and DAPI (blue), showing increased CD8^+^ T cell infiltration within the lesion of mice in B. Scale bars, as indicated. **h, i**, Quantification of CD8^+^ (h) and CD4^+^ (i) T cell density within lesion areas in g. **j**, Schematic model summarizing the proposed NR4A1-dependent mechanism. Genetic or pharmacologic inhibition of NR4A1 reduces NR4A1 binding to the *Nkg7* promoter, leading to up-regulation of cytotoxic effector molecules, NKG7 and possibly granzymes, which are responsible for enhanced CD8^+^ T cell cytotoxicity and improved control of *Mtb* infection. Data in b-i are representative of two independent experiments.

## Discussion

CD8^+^ T cells contribute to the control of *Mtb* infection in mice, macaques and humans ^45, 46, 47, 48^, however the relative importance and molecular mechanisms of CD8^+^ T cell-driven protective immunity in TB remain incompletely defined. We identified NR4A1 as a key transcriptional checkpoint, that restrains protective CD8^+^ T cell immunity in TB. Genetic ablation or pharmacological inhibition of NR4A1 altered CD8^+^ T cell states, promoting the expansion of activation and effector mechanisms while suppressing the exhaustion programs (Fig. 2 and 4). Together, these changes lead to enhanced *Mtb* control in vivo (Fig. 1 and 8). In line with the gene expression data, NR4A1-deficient mice showed increased frequency of lung Perforin^+^ and CD107a^+^ cytolytic CD8^+^ T cells along with decreased CTLA4^+^ and TIM3^+^ dysfunctional CD8^+^ T cells. Blocking TIM3 on T cells has been shown to enhance T cell function and decrease *Mtb* load in mice^10^. Notably, our data are consistent with the chronic infection models with lymphocytic choriomeningitis virus and Listeria monocytogenes, where pathogen-specific CD8^+^ T cells deficient in NR4A1 similarly displayed enhanced functionality, including increased cytokine production and cytolytic activity^18, 49^.

Importantly, our IF data in *Nr4a1*^⁻/⁻^ and DIM-C-treated WT mice demonstrated spatial inclusion of NR4A1-deficient CD8^+^ T cells in TB lesions (Fig. 6b-d and 8g-i). This may have greater significance in clinical settings since one of the factors limiting CD8^+^ T cell efficacy in TB is spatial segregation away from *Mtb*-infected macrophages^7, 8^. Cytolytic CD8^+^ T cells must contact target cells to eliminate *Mtb*^50^. The protective efficacy of NR4A1 inhibition aligned with the spatial transcriptomic results, which revealed enrichment of pathways related to cytotoxicity, natural killer cell-mediated killing, and interferon signaling specifically in the lung lesion T cell-rich regions (Fig. 6e-h). Collectively our findings demonstrate that ablation or inhibition of NR4A1 unleashes enhanced intralesional effector localization, accompanied by a spatially reinforced cytotoxic gene signature that drives more potent local immune control of *Mtb*.

Central to the NR4A1-mediated immunoregulatory circuit is NKG7, a granule-associated effector molecule transcriptionally repressed by NR4A1 via binding to the NR4A1-binding motifs (AAAGGTCA) in its promoter. We found NKG7 to be upregulated on CD8^+^ T cells when NR4A1 was inhibited in vivo and this effector molecule was responsible for the enhanced cytolytic activity of NR4A1-deficient CD8^+^ T cells against *Mtb* in vitro. In a tumor setting, NR4A1 binds directly to its motifs within promoter and enhancer regions of target genes, including *Ifng*, *Prf1*, and *Gzmb*, to repress their expression and promote tolerant/exhausted T cells^17, 18^. This repression is further enforced through interference with AP-1 activity, recruitment of chromatin modifiers, and establishment of a closed epigenetic landscape, as shown in genome-wide studies of exhausted CD8^+^ T cells in tumors and chronic infections^18^.

While our study provides compelling evidence that NR4A1 constrains protective CD8^+^ T cell responses during *Mtb* infection, the work has limitations. Although we focused on CD8^+^ T cells as the primary effector population affected by NR4A1 inhibition, NR4A1 is expressed in other cell types, including macrophages, CD4^+^ T cells and epithelial cells. We did not find evidence for enhanced activity of *Nr4a1*^-/-^ CD4^+^ T cells in the adoptive transfer or in vitro cytotoxicity assays, but we cannot exclude the possibility that non–T cell-intrinsic effects contribute to the observed phenotypes in global knockout or inhibitor-treated mice. Moreover, *Nr4a1 ^-/-^*mice exhibit spontaneous autoimmune features^31^ and NR4A1 shapes T cell development ^51^, these could act as confounding factors in our experiments. Future studies using multi-lineage conditional models or an inducible CD8^+^ T cell-specific *Nr4a1*-KO models will be critical to disentangle cell-type–specific roles.

In conclusion, our data in mouse models, accompanied by cross-species analyses, emphasize that reduced NR4A1 expression defines a transcriptionally reprogrammed CD8^+^ T cell state characterized by enhanced activation, cytolytic potential, and improved spatial access to *Mtb*-infected regions of the lung. These findings position NR4A1 as a tunable checkpoint for CD8^+^ T cell immunity in TB and provide a rationale for developing novel host-directed therapies for TB.

## Methods

### Study design

The objective of this study was to define the role of NR4A1 in host immunity to TB. We used *Nr4a1*^⁻/⁻^ mice infected with a low-dose aerosol infection of *Mtb* Erdman to assess the contribution of this transcription factor to disease progression. Adoptive transfer experiments were performed to determine the relative contribution of CD4^+^ versus CD8^+^ T cells from WT and *Nr4a1^⁻/⁻^* mice. To investigate the mechanisms underlying the *Nr4a1*^-/-^ dependent phenotype, we performed bulk RNA-seq, single cell RNA-seq, spatial transcriptomics, and other assays described below. All experiments were repeated at least twice independently, with the number of biological replicates mentioned in the figure legends. Sample sizes were based on previous experiments. Where applicable, outliers were excluded using GraphPad Prism.

### Mice

C57BL/6, *Nr4a1*^⁻/⁻^(KO), B6129PF2/J (WT), and *Rag1*^-/-^ mice were purchased from The Jackson Laboratory. All mice were maintained at UMass Chan Medical School under pathogen-free conditions. The health of the mice was monitored weekly by measuring body weight. Mice of both sexes between 6 and 10 weeks of age were used in accordance with a protocol approved by the Umass Chan Institutional Animal Care and Use Committee (IACUC). *Mtb*-infected mice were housed in biosafety level 3 (BSL3) facilities at the University of Massachusetts Chan Medical School.

### Aerosol infection in mice and bacterial quantification

Mice were infected via aerosol with *Mtb* Erdman using a Glas-Col Inhalation Exposure System (Terre Haute, IN), set to deliver 50-100 colony forming units (CFU) to the lungs, as confirmed by plating lung homogenates 24 hours after infection. At designated time points, mice were euthanized, and lungs and spleens were collected. Serial dilutions of tissue homogenates were plated on Middlebrook 7H11 agar plates for CFU enumeration.

### In vitro infections

For in vitro infections, we used two *Mtb* strains: H37Rv expressing a luciferase reporter (H37Rv-lux) and the clinical isolate CDC1551. Bacteria were cultured in Middlebrook 7H9 medium (Difco) to mid-logarithmic phase (OD₆₀₀ = 0.6–0.8). Thioglycollate-elicited peritoneal macrophages (TEPMs) were infected at a multiplicity of infection (MOI) of 1 for 3 hours, followed by three washes with fresh medium to remove extracellular bacteria. Infected macrophages were then incubated overnight at 37°C in a 5% CO₂ atmosphere. The following day (Day 1), purified CD8^+^ T cells were added to the cultures at an effector-to-target (E:T) ratio of 1:1. Bacterial burden was assessed by luminescence measurements (H37Rv-lux) and by plating on 7H11 plates (CDC1551) on Day 5 (post T cell addition).

### Generation of single-cell suspensions from the lung tissues

Lungs were isolated from mice and incubated in collagenase/DNAse for 30 min at 37 °C. Tissues were mechanically dissociated using a GentleMACS™ tissue dissociator (Miltenyi Biotec). The dissociated lung was passed through a 70-µm cell strainer to obtain a single-cell suspension, followed by red blood cells lysis and subsequent filtration through a 40-µm strainer to remove debris and aggregates.

### Lymphocyte isolation and adoptive transfer

Single-cell suspensions of homogenized spleens were prepared from naive mice. CD4^+^ and CD8^+^ T cells were purified by positive magnetic sorting using CD4 (L3T4) and CD8a (Ly-2) MicroBeads (Miltenyi Biotec), following the manufacturer’s protocol. The purified populations were stained and assessed for purity, which consistently ranged from 90% to 95%. For adoptive transfer experiments, 1 × 10⁶ purified CD4^+^ or CD8^+^ T cells were intravenously injected into *Rag1^⁻/⁻^* mice. Recipients were infected by aerosol with *Mtb* Erdman the following day.

### BCG vaccination

For vaccinations, BCG (SSI strain) was prepared at a concentration of 10⁶ CFU/mL in 0.04% Tween-80/PBS. Mice were administered with a single 100 μL subcutaneous injection. Sham-vaccinated controls received an equal volume of saline.

### Treatment with DIM-C-pPhCO2Me (DIM-C)

C57BL/6 mice were treated with the bis-indole-derived NR4A1 antagonist DIM-C-pPhCO2Me (Sigma) at a dose of 40mg/kg/day, administered every other day (3 times per week) for 21 days. Control mice were treated with vehicle-corn oil (Sigma).

### Flow cytometry

Lung single-cell suspensions were stained using standardized multicolor flow cytometry protocols. Cell viability was assessed using AmCyan-conjugated Live/Dead fixable dye (Invitrogen, L34966; 1:1000). Surface markers were stained in FACS buffer with antibodies against CD45 (Pacific Blue or APC-Cy7, clone 30-F11, BioLegend), CD3 (FITC, clone 145-2C11, BioLegend and BV785, clone 17A2, Biolegend), CD4 (PerCP-Cy5.5, clone GK1.5, BioLegend), CD8α (APC-Cy7, clone 53-6.7, BioLegend), CD44 (BV650, clone IM7, BioLegend), CD62L (PE-CF594, clone MEL-14, BD Pharmingen), NK1.1 (APC or Biotin, clone PK136, BioLegend), CD69 (BV605, clone H1.2F3, BioLegend),. Additional myeloid and tissue-resident markers included Ly6G (PE-CF594, clone 1A8), Ly6C (PE-Cy7, clone HK1.4), CD11b (BV605, clone M1/70), CD11c (PE, clone N418), CD43 (BUV737, clone S7), MHC II (BV650, clone M5/114.15.2), CD64 (BV421, clone X54-5/7.1), CD24 (PerCP-Cy5.5, clone M1/69), CD103 (APC, clone 2E7), CD90.2 (Biotin, clone 53-1.2), CD45R/B220 (Biotin, clone RA3-6B2), and MERTK (BV711, BioLegend). Immune checkpoint markers included PD-1 (BV421, clone RMP1-30, BD Biosciences), TIM-3 (BV711, clone B8.2C12, BioLegend), and CTLA-4 (PE/Fire 640, clone UC10-4B9, BioLegend). Streptavidin-BUV395 (BD Biosciences) was used for biotinylated secondary staining where appropriate. For intracellular cytokine and cytotoxic protein staining, cells were stimulated ex vivo using peptide pools or anti-CD3/CD28 in the presence of brefeldin A (BD Biosciences, 555028) and PE-Dazzle594–conjugated CD107a (BioLegend; clone 1D4B, 5 µg/mL). Following stimulation, cells were fixed and permeabilized using the BD Cytofix/Cytoperm kit and stained with antibodies against Granzyme A (PE, clone 3G8.5, BioLegend), Granzyme B (PerCP, Clone QA18A28, BioLegend), Perforin (APC, clone S16009A, BioLegend) and NKG7 (Cell signaling Technology, Clone E6S2A), all used at 1:50 dilutions. Data were acquired on a Cytek Aurora spectral cytometer and analyzed using FlowJo v10 (BD Biosciences).

### Histological analysis

For histology, lungs were inflated with 10% neutral-buffered formalin after perfusing with 1X Phosphate buffered saline. The multilobed right lung was immersion fixed in 10% buffered formalin for ≥24 hours. Lung tissue was processed, embedded in paraffin, sliced into 5 µm sections, and stained with hematoxylin and eosin (H&E) at the Morphology Core Facility, UMass Chan Medical School. The total lesion area and total lung area were quantified using ImageJ/Fiji and data were plotted using GraphPad Prism.

### Immunofluorescence staining, Imaging and analysis

Formalin fixed paraffin-embedded (FFPE) lung sections were used for IF staining. Slides were deparaffinized, rehydrated, and subjected to antigen retrieval, followed by overnight incubation with fluorophore conjugated primary antibodies at 4°C. The primary antibodies used were anti-CD4 (Abcam EPR19514) and anti-CD8 (Abcam EPR21769). Images were acquired using a TissueFAXS iQ tissue cytometer equipped with a Hamamatsu ORCA Fusion BT camera, and quantification was performed in QuPath software^52^. For each mouse, immunofluorescence imaging was performed across all lobes of the multilobed right lung. All lesion-containing regions within these lobes were identified and imaged, resulting in variable numbers of regions per mouse due to higher lesion burden in WT and fewer lesions in *Nr4a1*^⁻/⁻^ mice. Lesions were manually annotated in QuPath, and CD4^+^/CD8^+^ T cells were classified based on clear full-rim membrane fluorescence. Lesion cores were manually annotated based on the central macrophage-dense or necrotic region of the lesion, clearly distinct from the peripheral lymphocyte enriched rim. For each mouse, total positive cell counts were summed across all lesion regions and normalized to the total annotated lesion area.

### Quantitative real-time PCR (qPCR)

Total RNA was extracted from cells using the TRIzol reagent (Thermo Fisher Scientific) following the manufacturer’s instructions. RNA quality and concentration were assessed using a NanoDrop 2000 spectrophotometer (Thermo Fisher Scientific). Complementary DNA (cDNA) was synthesized from 1 µg total RNA using the High-Capacity cDNA Reverse Transcription kit (Applied Biosystems). Quantitative PCR (qPCR) was performed using AzuraQuant Green Fast qPCR Mix (Azura Genomics) on a CFX Opus 96 Real-Time PCR System (BioRad). Relative mRNA expression was calculated using the 2⁻^ΔΔCt^ method, with β-actin used as the endogenous control. Primer sequences for all target genes are provided in Supplementary Table 6. All reactions were performed in technical triplicates, and data represent at least three independent biological replicates.

### Luminex Bioplex assay

Multiplex cytokine/chemokine analysis was performed using the Bio-Plex Pro Assay platform (Bio-Rad, USA), following the manufacturer’s instructions. Briefly, filtered lung tissue homogenates were collected and stored at -80°C until analysis. Samples were thawed on ice and diluted before incubation with magnetic beads coated with target-specific capture antibodies from ProcartaPlexTM Mouse Cytokine & Chemokine Convenience Panel 1A, 36plex (ThermoFisher; Cat no. EPXR360-26092-901). Fluorescence intensity was measured using a Bio-Plex 200 System (Bio-Rad), and data were analyzed using Bio-Plex Manager software. All samples were run in triplicates and results were expressed as pg/mL.

### Flex scRNA-seq library preparation and sequencing

CD90.2 T cells from *Mtb*-infected and uninfected WT and *Nr4a1*^⁻/⁻^ mice were sorted and fixed using the Chromium Next-GEM Single-Cell Fixed RNA Sample Preparation Kit (10x Genomics). Four fixed samples were multiplexed in a single reaction by hybridizing four distinct barcoded mouse transcriptome probe sets from the Chromium Fixed RNA Kit (10x Genomics). For each pool of four samples, 20,000 cells were targeted for capture on the Chromium iX instrument, followed by library generation following the manufacturer’s protocol for 10x Genomics Flex Gene Expression. The resulting libraries were sequenced on an Illumina NovaSeq X Plus system, targeting 25,000 read pairs per cell.

### scRNA-seq data analysis

Raw Flex scRNA-seq reads in FASTQ format were aligned to the mouse reference genome (mm10) using Cell Ranger multi (v7.1.0). The resulting demultiplexed outputs from eight libraries were then imported into Seurat (v4.4.0) in R (v4.4.2) for downstream analysis. Quality control (QC) was conducted on each library to filter out low-quality cells based on the following criteria: (1) cells with fewer than 300 or greater than 97th percentile of detected genes (nFeature_RNA), (2) cells with fewer than 500 or greater than the 97th percentile of total RNA molecules (nCount_RNA),and (3) cells with over 5% of reads mapped to mitochondrial genes. After normalization, DoubletFinder (v2.0.3) was used to identify and exclude doublets from each library. The eight Seurat objects were then integrated into a single dataset. Dimensionality reduction was performed using PCA followed by UMAP, and unsupervised clustering was conducted using a graph-based algorithm with a resolution of 1.2. To achieve finer resolution of CD8^+^ T cells, clusters expressing *Cd3d* and *Cd8* genes were isolated and re-clustered at the same resolution. T cells expressing *Cd4* and *Trdc* genes were removed prior to downstream analysis. The remaining CD8^+^ T cell subtypes were annotated based on their gene expression profiles, including (1) naïve T cells (T_n_) expressing *Ccr7* and *Sell*, (2) effector memory T cells (T_em_) expressing high levels of *Gzmk*, (3) regulatory T cells (T_reg_) highly expressing *Il2ra*, (4) tissue-resident memory T cells (T_rm_) expressing *Itgae* and *Cdh1*, (5) metabolically-active T cells highly expressing mitochondrial genes such as *mt-Co2* and *mt-Atp6*, (6) effector T cells (T_e_) highly expressing *Gzma*, (7) exhausted T cells (T_ex_) highly expressing *Tox* and *Pdcd1*, and (8) proliferating T cells (T_prolif_) expressing *Mki67*. Notably, *S100a4* was highly expressed across all non-naïve T cell subsets.

Differential expression analysis between *Nr4a1*^⁻/⁻^-Inf (KO-Inf) and WT-Inf cells was performed for each CD8^+^ T cell subtype using the *FindMarkers* function from Seurat v5. The parameters were set to *logfc.threshold=0* and *min.pct=0* to include all genes in the analysis. Genes with an adjusted *P* value <0.05 were considered DEGs. For pseudobulk DEG analysis, all CD8+ T cell subtypes were aggregated to form a combined population prior to comparison. Pathway enrichment analysis of DEGs was performed using the clusterProfiler R package (v4.14.6). All genes (n = 34,089) present in the integrated Seurat object were used as the background gene set. Gene symbols were first converted to ENTREZ IDs using *bitr* function from clusterProfiler and org.Mm.eg.db annotation package (v3.20.0) for the mouse genome. GO over-representation analysis was carried out using *compareCluster* function, with the following parameters: *fun=’enrichGO’*, *ont=’BP’*, *pvalueCutoff=0.05*, and *qvalueCutoff=0.10*. Redundant GO terms were manually removed, and the final enrichment results were visualized using the *dotplot* function from clusterProfiler.

### Spatial transcriptomics using GeoMx Digital Spatial Profiler

Serial sections (5 μm) from FFPE blocks were taken from selected samples, to prepare slides for RNA hybridization and profiling. Slides were baked at 60 °C for 30 minutes prior to deparaffinization, then subjected to antigen retrieval (EDTA-based pH 9, 15 minutes, >99°C). Enzymatic exposure of RNA targets (1μg/mL Proteinase K, 15 minutes, 37°C) was performed, then slides were post-fixed with 10% Neutral Buffered Formalin and washed. Slides were then hybridized overnight with GeoMx Mouse Whole Transcriptome Atlas (Nanostring) at 37 °C in a humidified chamber. After stringent washing (4X SSC + 100% Formamide, 1:1), slides were blocked with Buffer W (GeoMx RNA Slide Prep Kit) for 30 min at room temperature, then stained with the following in Buffer W for 1 hour at room temperature: SYTO13 (1:10000, Invitrogen), CD3-A647 (1:100, Biorad), CD45-A594 (1:20, Cell Signaling), CD11b-A532 (1:40, Novus). Slides were washed twice with 2X SSC before loading onto the GeoMx DSP instrument for scanning, ROI selection and segmentation. Each ROI was segmented into compartments for collection in a sequential manner; Lymphocytes (CD3^+^), Myeloid (CD11b^+^), Immune (CD45^+^) and Stroma (-residual tissue). Each segment was then eluted and placed in a well in a 96-well collection plate. Eluted probes were amplified using GeoMx Seq Code Pack (Nanostring) and pooled according to the manufacturer’s protocol. The pooled libraries were sequenced on an Illumina HiSeq X lane with 5% PhiX spike-in, targeting 100 reads per square micron of total segment area.

### Spatial transcriptomics data analysis

Sequencing data were processed in the NanoString (now Bruker) GeoMx NGS Pipeline to generate DCC files, which were analyzed with R packages GeomxTools (v3.10.0) and NanoStringNCTools (v1.14.0). Areas of illumination (AOIs) were included if they met quality-control criteria: >1,000 raw reads, >80% trimmed reads, >80% stitched reads, >75% aligned reads, sequencing saturation >50%, segment area >1,000, >20 estimated nuclei, >1 negative control counts, and <9,000 counts in the NTC well; one AOI was excluded for low read count. Probe-level QC was performed using *setBioProbeQCFlags* function with default parameters (*minProbeRatio=0.1, percetFailGrubbs=20, removeLocalOutliers=True*), and the gene-level count matrix were derived by computing the geometric mean of read counts mapped to multiple probes targeting the same gene. Genes were retained if detected above the limit of quantification (LOQ) in >5% of AOIs, and data were normalized using Q3 normalization based on the top 25% most highly expressed targets. Differential expression analysis between *Nr4a1*^⁻/⁻^ and WT groups in core CD3^+^ cells was performed using *mixedModelDE* function by specifying section ID (i.e mouse ID) as a random effect; genes with log2FC>0.5 and nominal P<0.05 were considered significant.

### Bulk RNA-seq library preparation and sequencing

Total RNA was extracted from mouse spleen CD8^+^ T cells lysed in TRIzol (Thermo Fisher), using acid guanidinium thiocyanate-phenol-chloroform extraction followed by a Qiagen RNeasy Micro clean-up procedure. RNA was analyzed on Agilent Bioanalyser for quality assessment. cDNA libraries were prepared using 2 ng of total RNA using the Smart-seq2 protocol ^53^ with the following modifications: 1. Addition of 20 µM TSO; 2. Use of 200 pg cDNA with 1/5 reaction of Illumina Nextera XT kit. The length distribution of the cDNA libraries was monitored using a DNA High Sensitivity Reagent Kit on the Perkin Elmer Labchip. All samples were subjected to 2×151 sequencing on an Illumina NovaSeq 6000 system targeting 15 million read pairs per sample. GO over-representation analysis was carried out using *compareCluster* function as described in scRNA-seq data analysis section above.

### Bulk RNA-seq data analysis

Raw sequencing reads in FASTQ format were aligned to the mouse reference genome GRCm39 using the STAR aligner (v2.7.10b). Gene-level counts were obtained with the featureCounts function from the Rsubread package (v2.8.2) in R. These counts were then transformed into log₂ RPKM values using the edgeR package (v4.4.2). PCA analysis was then performed on log₂ RPKM values using FactoMineR package (version 2.11). Differential gene expression analysis was performed using the DESeq2 package (v1.46.0), with p-values from the Wald test adjusted for multiple comparisons via the Benjamini-Hochberg procedure. Genes were considered significantly differentially expressed if they had a false discovery rate (FDR) below 0.05 and a log₂ fold change greater than 1.5 or less than -1.5. Time-course DEGs between *Nr4a1*^⁻/⁻^ and WT mice were identified at 2, 4, and 8 weeks using uninfected mice as baseline, with the main terms being genotype and time and interaction term being genotype:time.

### Analysis of publicly available macaque and human datasets

Single-cell transcriptomics data of *Mtb*-infected macaques ^41^ was obtained from Broad Single-Cell Portal (accession code: SCP2689). Raw gene expression data stored in an AnnData object was extracted using scanpy package (v1.11.1) in Python (v3.10.12) and loaded into R using *open_matrix_anndata_hdf5* function from the BPCells package (v0.3.0). A Seurat object was created from this raw expression data, incorporating associated metadata and UMAP coordinates. CD8^+^ T cells from the naïve cohort, including Tc17, T_Eff_, GZMK^hi^ T_EM/PEX_, and T_EMRA_ subsets as described by Bromley et al, were isolated, and dimensional reduction was performed on this subset. These cells were classified into CD8^+^NR4A1^lo^ (T_Eff_, GZMK^hi^ T_EM/PEX_, T_EMRA_) and CD8^+^NR4A1^hi^ (Tc17) populations based on relative NR4A1 expression levels.

Human lung single-cell transcriptomics data from active TB patients ^42^ was obtained from the GEO database (accession code: GSE192483). The filtered_feature_bc_matrix.h5 files for samples SP019H, SP019L, SP020L, SP021H, SP021L, SP023H, SP023L, SP024H, SP024L, SP025H, and SP025L were processed using the Seurat v5 package in R. QC, filtering, and doublet removal were performed as described in the “Flex scRNA-seq data analysis” section above. Data integration was conducted using the *IntegrateLayers* function in Seurat v5. After dimensional reduction, clustering was carried out at a resolution of 1.0, followed by sub-clustering of CD8^+^CD3^+^ cells at a resolution of 0.8. One sub-cluster (cluster 3), identified as TRDV1^+^ and likely representing contaminating γδ T cells, was removed. The remaining eight clusters were grouped into CD8^+^NR4A1^lo^ (clusters 0, 2, 4, 6) and CD8^+^NR4A1^hi^ (clusters 1, 5, 7, 8) populations based on NR4A1 expression. Only FDG^hi^ samples were included in the downstream differential expression analysis.

For both datasets, DEG analysis between CD8^+^NR4A1^lo^ and CD8^+^NR4A1^hi^ populations was performed using *FindMarkers* function from Seurat v5. Genes with adjusted P<0.05 were considered significant. Pathway enrichment analysis was carried out using Enrichr with the BioPlanet_2019 library.

### siRNA knockdown of *Nkg7* in mouse CD8^+^ T cells

CD8^+^ T cells were purified using magnetic sorting by positive selection with CD8a^+^ (Ly-2) MicroBeads (Miltenyi Biotec), following the manufacturer’s protocol. For siRNA transfection, cells were seeded at 1 × 10^6^ cells per well in a 96 well plate. Accell mouse *Nkg7* siRNA set (Horizon Discovery/Dharmacon; A-063689-15-0010) consisting of four individual Accell siRNAs targeting *Nkg7* was used for gene knockdown. A non-targeting Accell siRNA control pool (D-001910-10-05) was used as control. Briefly, siRNA was reconstituted in nuclease-free water to a 100 μM stock concentration and diluted in Accell Delivery Media to a final concentration of 1 μM. The culture medium was replaced with siRNA-containing Accell Delivery Media, and cells were incubated at 37°C with 5% CO₂ for 24-72 hours to allow siRNA uptake and gene silencing; as per the manufacturer’s protocol. Gene knockdown efficiency was assessed by quantitative PCR (qPCR) for mRNA expression.

### Chromatin Immunoprecipitation (ChIP)–qPCR

Splenic CD8^+^ T cells from C57BL/6 mice were isolated by magnetic sorting (Miltenyi Biotec). For each ChIP reaction, 1.5 × 10⁷ CD8^+^ T cells were used. Cells were stimulated with phorbol 12-myristate 13-acetate (PMA, 20 ng/ml) and ionomycin (1 µg/ml) for 6 h at 37 °C prior to chromatin preparation. ChIP assays were performed using the ChIP-IT PBMC Kit (Active Motif, 53042) following the manufacturer’s instructions. Chromatin was immunoprecipitated using anti-NR4A1 (NGFI-Bα/Nur77) antibody (5 µg per reaction, Novus Biologicals, # NB100-56745) or normal IgG control. The recovered DNA was analyzed by quantitative PCR using *Nkg7* promoter specific primers and AzuraQuant^TM^ GreenFast qPCR Mix (AZ-2105) on a CFX Opus 96 Real Time PCR System (Biorad). Raw Ct values from ChIP and input DNA were used to calculate enrichment relative to input (% input) using the formula: % Input = 100 × ^2[Ct(Input) – Ct(ChIP)]^. For comparison across samples, enrichment was normalized to IgG controls and data are presented as percent input.

### Data visualization and Statistical analysis

All dot plots and bar graphs were generated using GraphPad Prism (v9.5.1), with statistical analyses detailed in the respective figure legends. UMAPs, feature plots, violin plots, and gene expression bubble plots for single-cell and spatial transcriptomics data were produced using the Seurat package in R. The stacked violin plot was generated with the scCustomize package (v3.0.1). Volcano plots were created using the EnhancedVolcano package (v1.24.0), and heatmaps were generated using the pheatmap package (v1.0.12). Schematic diagrams illustrating the experimental workflows were created using BioRender (https://www.biorender.com).

## Acknowledgement

We thank Heather Farineau for technical assistance; the Morphology Core Facility at UMass Chan Medical School (UMMS) for H&E staining; and the Core BSL-3 Facility for access to equipment and infrastructure. We also acknowledge Drs. Nuria Martinez and Christina Baer for support with preliminary mouse experiments and immunofluorescence imaging, respectively. We are grateful to Dr. Batuhan Yenilmez for assistance with siRNA experiment. This work was supported by grants to H.K. and A.S. from The National Heart, Lung, and Blood Institute (R01HL153162 and R01HL152078) and The National Institute of Allergy and Infectious Diseases (R21AI185410) of The National Institutes of Health (NIH). A.S. is supported by A*STAR ID Labs, BMRC CRF and Singapore’s Trident grant. NIH funds ($545,150) accounted for 82% of the total project costs, while other funds ($120,000) accounted for the remaining 20%. The content is solely the responsibility of the authors and does not necessarily represent the official views of the NIH.

## Author contribution

S.F. designed and performed experiments, collected and interpreted data, and generated the initial draft of the manuscript. S.S., H.R. and A.B. carried out spatial transcriptomic investigation. A.T., S.Foo and S.W.H. generated bulk and scRNA-seq data. L.S., B.S., C.J. and M.J. assisted with mouse experiments and data acquisition. Y.C. performed bioinformatics analyses and contributed to the initial draft of the manuscript. All authors contributed to the discussion. A.S. and H.K. conceived and designed the study, acquired funding, supervised the research, interpreted data, and revised the manuscript.

## Inclusion & Ethics Statement

The authors affirm that this study was conducted in accordance with ethical principles, and that inclusion, diversity, and global research equity were considered in the design, execution, and authorship of this work.

## Data Availability Statement

All data supporting the findings of this study are available within the article, its Supplementary Information, or from the corresponding authors upon reasonable request. The bulk and single-cell RNA-seq data generated in this study has been deposited in the GEO repository (GSE313371 and GSE313372, respectively). Spatial transcriptomics datasets will be deposited upon acceptance. Publicly available macaque and human single-cell datasets used in this study are cited and linked in the Methods.

## Code Availability Statement

No custom code was generated for this study. All analyses were performed using publicly available software and packages, including Cell Ranger, Seurat, DoubletFinder, clusterProfiler, EnhancedVolcano, FlowJo, GraphPad Prism, GeomxTools, and standard R packages. Scripts used for data processing and visualization are available from the corresponding authors upon reasonable request

## Legends Supplementary tables

**Supplementary Table 1** DEGs identified from the time-course RNA-seq analysis of splenic CD8^+^ T cells from WT and *Nr4a1^⁻/⁻^* mice.

**Supplementary Table 2** DEGs in splenic CD8^+^ T cells from WT and *Nr4a1^⁻/⁻^* mice at 4 and 8 wk after *Mtb* infection (Fig. 2d). Includes cluster assignments for 4 wk DEGs (clusters i–x; Extended Data Fig. 2b) and 8 wk DEGs (clusters A–J; Fig. 2e).

**Supplementary Table 3** DEGs from scRNA-seq analysis of lung CD8^+^ T cell subsets from WT and *Nr4a1^⁻/⁻^* mice at 4 wk. Includes gene lists for *Gzmk^+^* effector-memory (cluster g) and *Gzma^+^* effector (cluster m) populations (Fig. 4e).

**Supplementary Table 4** DEGs from GeoMx DSP whole-transcriptome analysis of CD3^+^ T cell ROIs in lung lesions at 4 wk.

**Supplementary Table 5** DEGs stratified by NR4A1 expression state (NR4A1^lo^ versus NR4A1^hi^) in CD8^+^ T cells from macaques and human TB datasets

**Supplementary Table 6** Sequences of primers

## Notes

### Competing Interest Statement

The authors have declared no competing interest.

