## Supplementary Figures and Tables for "NR4A1 limits CD8^+^ T Cell effector responses and protection in tuberculosis"

Extended Data Figure 1

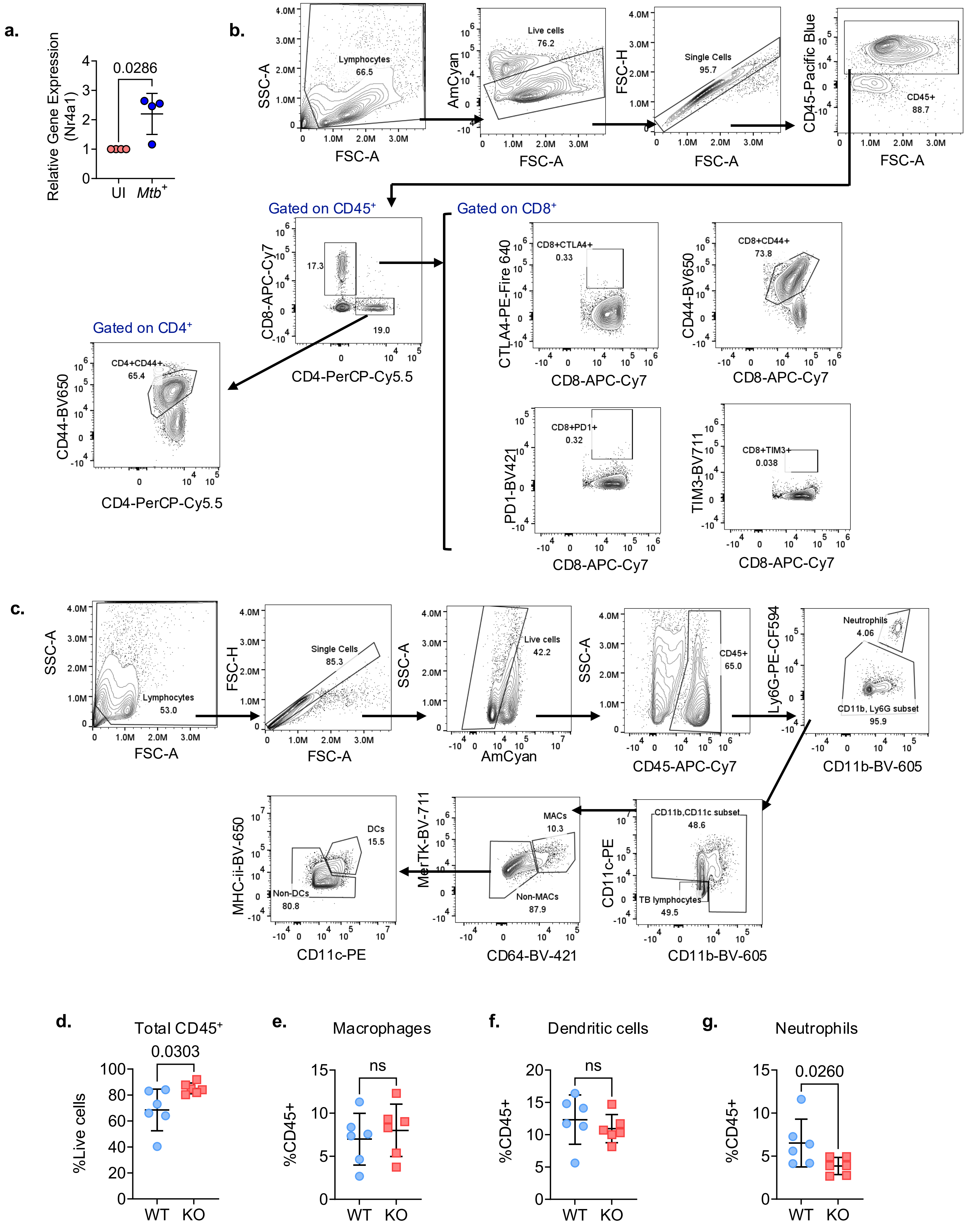

**Extended Data Fig. 1 Flow-cytometric gating and quantification of T cell and myeloid subsets in *Nr4a1*-deficient versus WT mice lungs.**

**a,** Lung *Nr4a1* mRNA levels in uninfected (UI) and *Mtb*-infected mice, shown as relative expression (qPCR). *P* value, two-tailed Mann–Whitney. Each dot represents one mouse. Data is mean ± s.d. **b,** Flow cytometry gating for lung T cells using activation and exhaustion markers. **c,** Gating strategy for analyzing myeloid cells in the lung. **d,** Percentage of $CD45^+$ T cells among live lung cells in WT and *Nr4a1*$^{-/-}$ mice. **e–g,** Macrophages **(e),** dendritic cells **(f)** and neutrophils **(g)** as a percentage of $CD45^+$ lung leukocytes. Each dot is a mouse; horizontal bar indicates mean ± s.d. P value, two-tailed Mann–Whitney test.

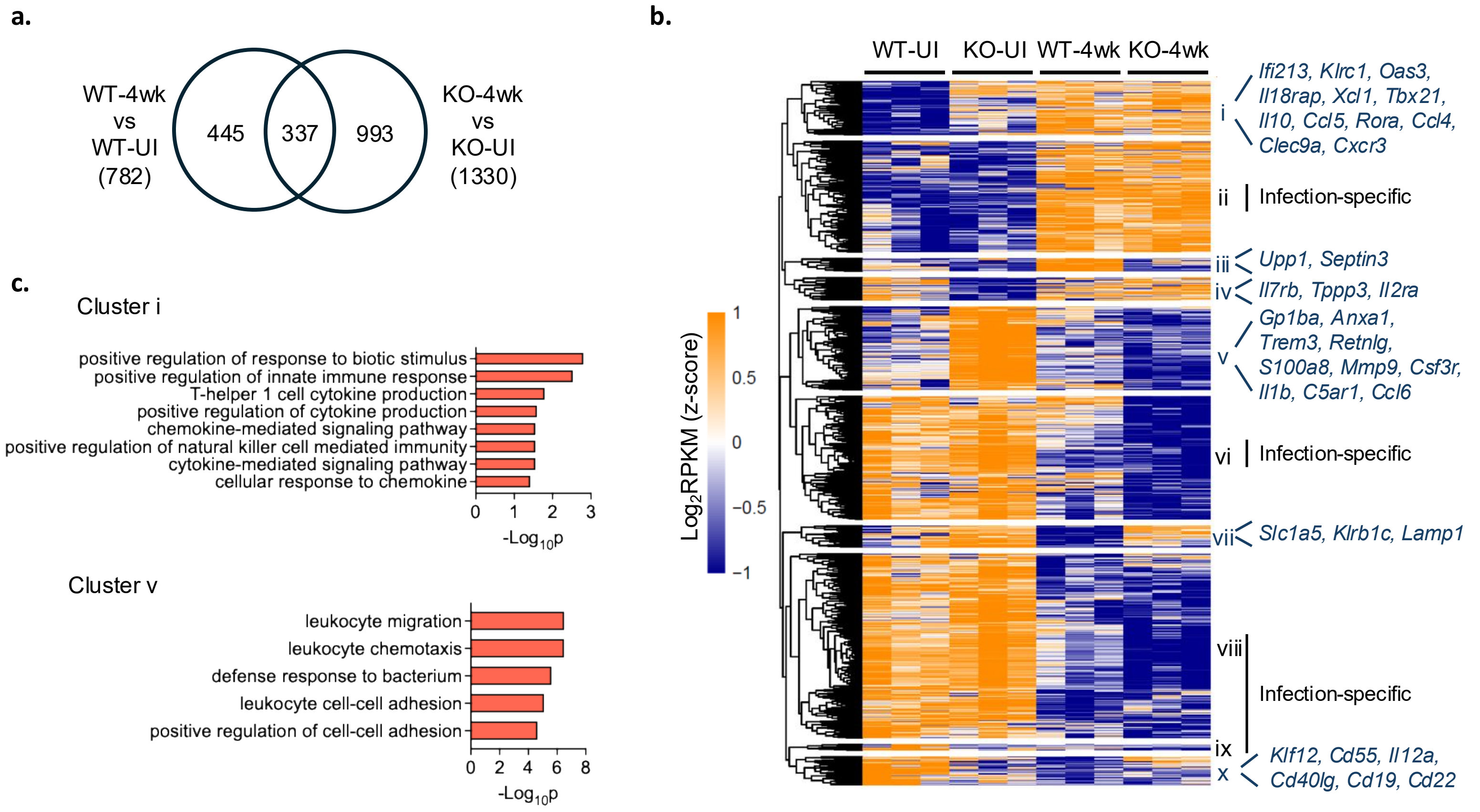


**Extended Data Fig. 2 Bulk RNA-seq of spleen CD8⁺ T cells from uninfected and infected (4 wk) Nr4a1$^{-/-}$ and WT mice.**

**a**, Venn diagram showing the overlap of DEGs between 4 wk and uninfected (UI) CD8$^+$ T cells from WT and *Nr4a1*$^{-/-}$ mice. **b**, Heatmap of the union of 4 wk vs UI DEGs (*n* = 1,775) identified in panel a. $Log_2$ RPKM values were *z*-score–normalized across samples. Genes were grouped into ten clusters (cluster i to x) using *k*-means clustering. Representative genes for clusters not classified as infection-specific are labeled on the right. **c,** GO biological processes among clusters i and v from panel b. Enriched biological processes include pathways related to cytokine-mediated signaling, natural-killer-like effector activity, leukocyte chemotaxis, and antibacterial defense. GO terms shown represent non-redundant categories with the lowest adjusted *P* values.

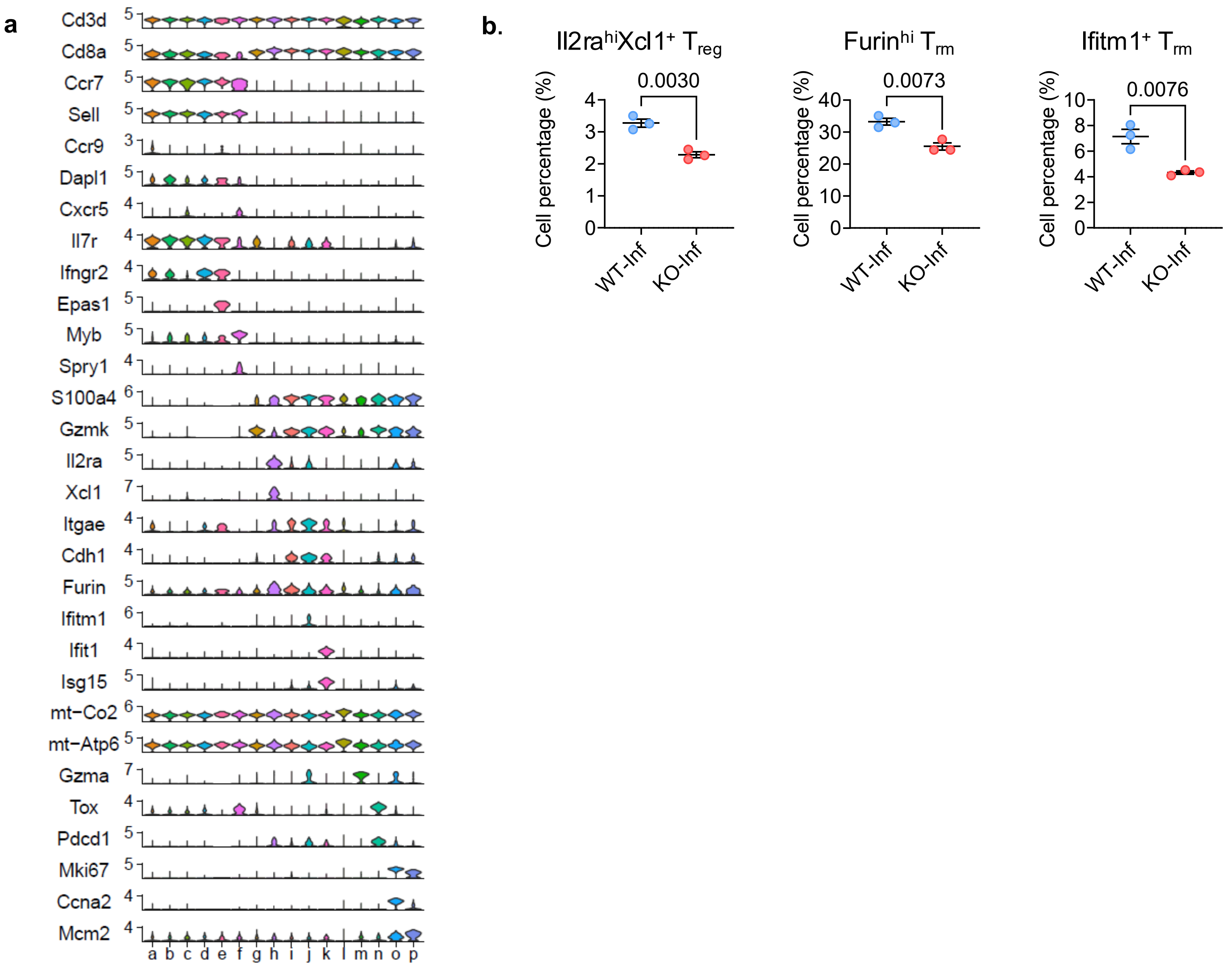


**Extended Data Fig. 3 Marker expression and subset frequencies from scRNA-seq data define CD8$^+$ T cell states in WT and *Nr4a1*$^{-/-}$ lungs.**

**a**, Expression of markers used to characterize CD8$^+$ T cell subtypes (see Methods for gene lists and normalization). **b**, Percentage of Il2ra$^{hi}$ Xcl1$^+$ $T_m$, Furin$^{hi}$ $T_{rm}$ and Ifitm1$^+$ $T_{rm}$ in infected lungs (WT versus *Nr4a1*$^{-/-}$); unpaired two-sided *t*-test; *P* values labeled.

Extended Data Figure 4

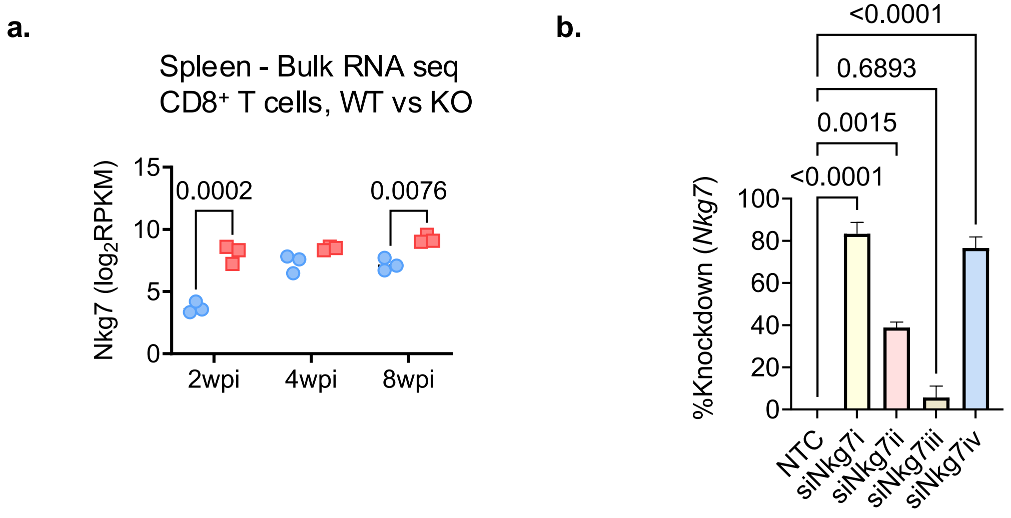


**Extended Data Fig. 4 *Nkg7* expression and its knockdown in CD8$^+$ T cells.**

**a,** Bulk RNA-seq analysis showing *Nkg7* expression ($\log_2$ RPKM) in splenic CD8$^+$ T cells from WT and *Nr4a1*$^{-/-}$ mice at 2, 4, and 8 wk of infection. *P* value, unpaired two-tailed *t* test. **b**, Efficiency of *Nkg7* knockdown in *Nr4a1*$^{-/-}$ CD8$^+$ T cells using four independent siRNAs, as measured by RT-qPCR 72 h post-transfection. Data mean ± s.d.; one-way ANOVA.

Extended Data Figure 5

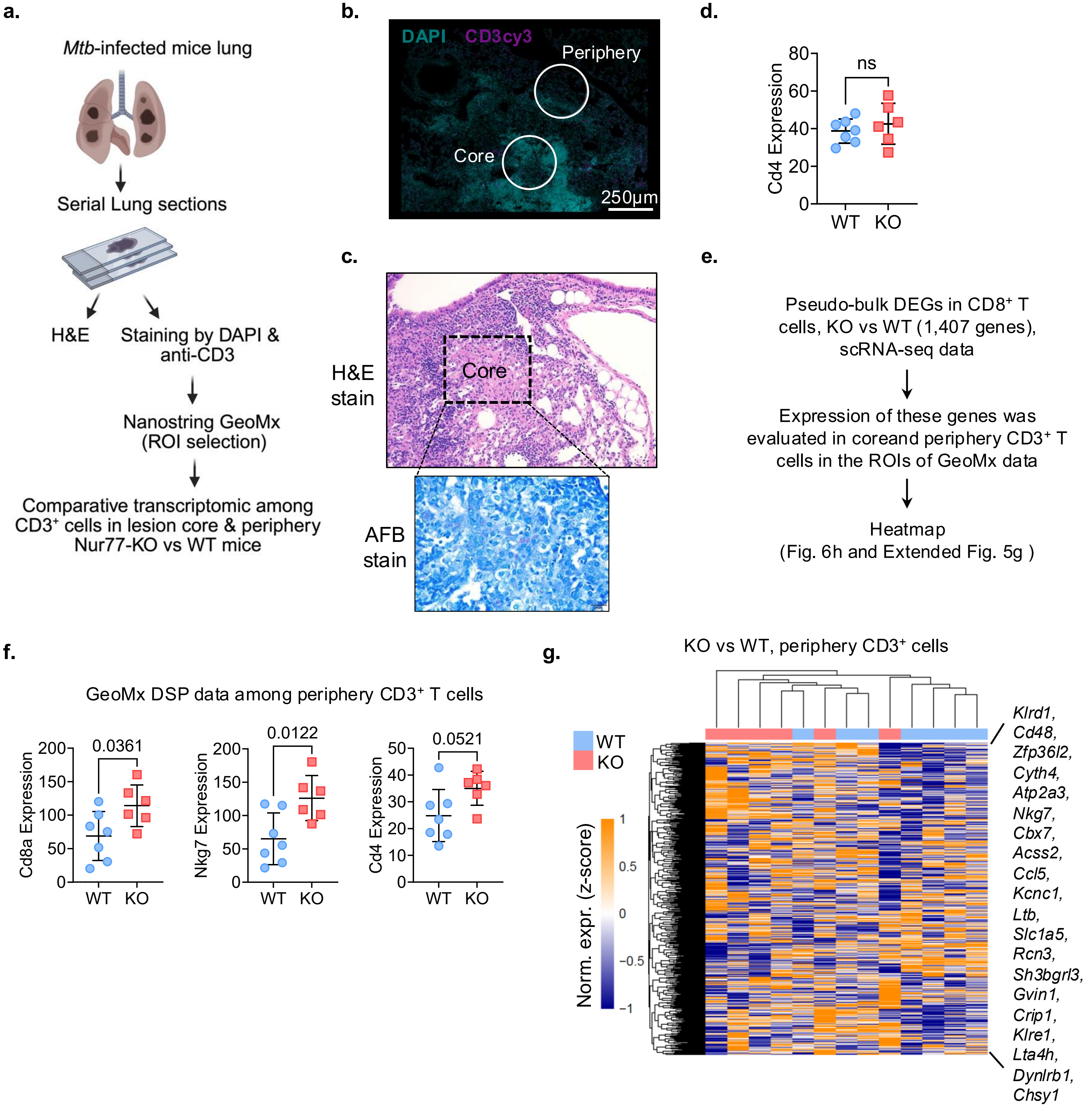

**Extended Data Fig. 5 Spatial transcriptomic workflow and key readouts in *Mtb*–infected mice lungs.**

**a,** Schematic of GeoMx spatial transcriptomics: serial lung sections; H&E and antibody staining (DAPI, CD3, CD11b) to guide ROI selection; comparative transcriptomics of $CD3^+$ and $CD11b^+$ cells in lesion core versus periphery, *Nr4a1*$^{-/-}$ versus WT. **b,** Immunofluorescence image showing ROIs in lesion core and periphery defined by DAPI and CD3 (Cy3) signal; scale bar, 250 µm. **c,** Histology of infected lung sections: H&E (top) and acid-fast bacilli (AFB) stain (bottom). **d,** *Cd4* expression (normalized counts) in $CD3^+$ ROIs from WT and *Nr4a1*$^{-/-}$ (KO) lungs (*P* value, two-sided Mann–Whitney test). **e,** Schematic for the heat map generation by taking DEGs from $CD8^+$ T cell scRNA-seq data and finding their gene expression in periphery or core $CD3^+$ ROIs of spatial transcriptomic data. **f,** Expression levels of *Cd8a*, *Nkg7*, and *Cd4* in peripheral $CD3^+$ T cells from WT and *Nr4a1*$^{-/-}$ mice. *P* values, unpaired two-tailed *t* tests. **g,** Heatmap of the pseudo-bulk DEGs identified from total lung $CD8^+$ T cell scRNA-seq (Fig. 5b), evaluated in periphery $CD3^+$ ROIs from GeoMx spatial transcriptomics of *Mtb*-infected lungs. Distinct clustering of WT (blue) and *Nr4a1*$^{-/-}$ (red) samples was not observed among periphery $CD3^+$ ROIs. Representative cytotoxic and activation genes are indicated on right side. See Methods for DEG definition, normalization and clustering parameters.

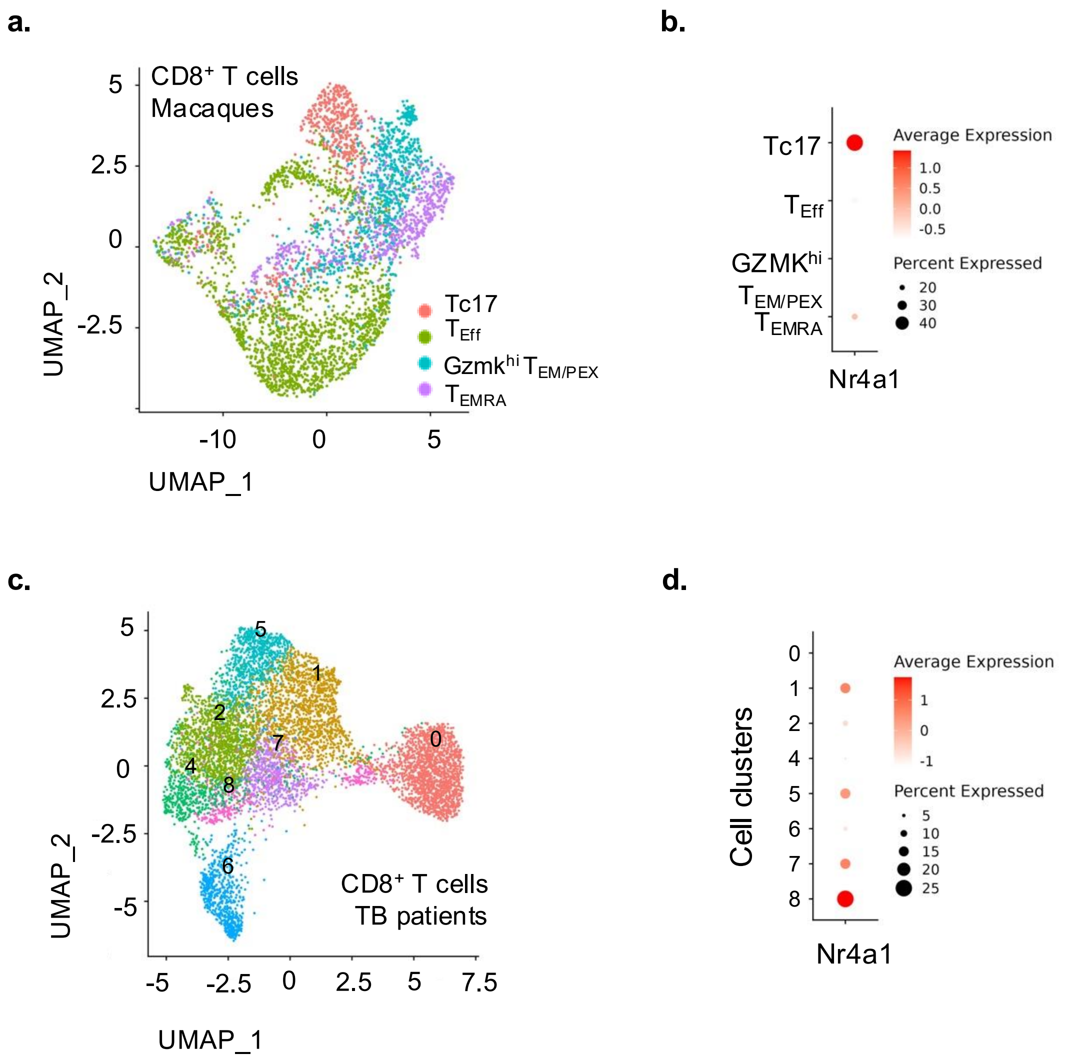


**Extended Data Fig. 6 *Nr4a1* expression across lung CD8⁺ T cell states in macaque and humans with TB, analysis of published scRNA-seq dataset.**

**a,** UMAP of four lung CD8⁺ T cell subtypes in *Mtb*-infected macaques. **b,** Bubble plot summarizing *Nr4a1* expression across the four lung CD8⁺ T cell subtypes in **a** (dot size, percent expressing; color, average normalized expression). **c,** UMAP of eight CD8⁺ T cell clusters in the lungs of active TB patients. **d,** Bubble plot summarizing *Nr4a1* expression across the eight clusters i**n c** (dot size, percent expressing; color, average normalized expression). Dataset sources and accessions are listed in Methods.
